# Cell Cycle Sensing Shapes Human T Cell Fate and Exhaustion Programs

**DOI:** 10.64898/2026.06.11.731737

**Authors:** Meelad Amouzgar, Tara Murty, Patricia Favaro, Elena Sotillo, Trevor Bruce, Daniel Ho, Avery J. Lam, Robert Tibshirani, Crystal L. Mackall, Sean C. Bendall

## Abstract

Cell cycle (CC) dynamics are reflected in diverse T cell processes such as TCR activation, expansion, contraction, differentiation, senescence, anergy, and exhaustion; linking CC behaviors to functional and dysfunctional T cell states in development and disease. Progression through CC checkpoints is also tightly linked to cell fate decisions across development. Yet, much remains unknown about the connection between CC sensing and T cell differentiation programs. To disentangle the relationship across T cell state, time-since-activation, receptor signaling, division, and CC, we leverage high-throughput single-cell mass cytometry for parallel measurement of these diverse biological states. By modulating CC progression and receptor signaling with inhibitors as well as tonic signaling Chimeric Antigen Receptor (CAR) models of T cell exhaustion, we reveal that earlier G1/S CC programs crosstalk with receptor signaling to control T cell fate, and that exhaustion programs are downstream to aberrant, S-G2 phase CC arrest signatures in tonic CAR signaling *in vitro*, *in situ, and in vivo* across human cancers in association with CD8 T-lymphocyte dysfunction.

## Introduction

Cell cycle (CC) progression and division are tightly linked to cell fate decisions across the hematopoietic system.^1–5^ In T cells—central players of adaptive immunity—recognition of presented antigen and co-stimulation triggers rapid CC entry, inducing a massive, temporally coordinated wave of proliferation that drives expansion into varying functional phenotypes of effector cells followed by a contraction that favors the persistence of long-lived memory populations.^6–10^ This tightly controlled expansion-contraction is paired with loss of effector functions in T cell populations to protect against excessive immune activation and autoimmunity, contextualizing that T cell function and exhaustion (i.e., dysfunction) are evolved mechanisms of protection as a safeguard from natural T cell processes.^11–18^ Within this context, CC dynamics are both a regulator and a reflection of T cell processes such as TCR activation, expansion, senescence, anergy, and exhaustion, linking CC behavior to the emergence of functional and dysfunctional T cell states in development, immunity, and disease.^19–22^

Studies looking at T cell (dys)function largely focus on static snapshots of time, where upstream and downstream biology of cellular states are inferred through measurement of the gain or loss of T cell related molecules and effector functions. One such example is T cell exhaustion, characterized by loss of effector abilities (i.e., reduced cytotoxicity and cytokine production), expression of activation & inhibitory receptors, and importantly, impaired proliferation.^13,23,24^ This correlates cell cycle aberrancy with T cell dysfunction, but developmental biology teaches that CC programs are often intimately linked in controlling cell differentiation and function^1,3,4,22,25^. In T cells, cell cycle–arrested states are linked to T cell fate with early CDK4/6 inhibition promoting memory formation, enhanced effector function, and metabolic switches, and evidence suggests that T cell fate can be locked-in as early as the first division^20,26,27^. Furthermore, successful T cell genetic engineering strategies that enhance key T cell programs like effector function and proliferation often involve targeting powerful CC regulators, such as the Mediator cyclin-dependent kinase module (i.e., MED12 subunit), components of the Activator Protein-1 complex (i.e., c-Jun), and IKZF3.^28–31^

Despite TCR signaling and CC programs being consequential for T cell fate and function, a detailed map of cell cycle progression and division dynamics coupled to T cell state has not been elucidated. This is partially because capturing the early temporal coordination of proliferation, cell cycle states, and T cell fate decisions in primary human T cells requires a high-dimensional, parallel profiling approach—beyond what conventional static snapshots can provide. Additionally, T cell (dys)function studies narrowly study CC state as a correlative consequence of T cell state rather than a partner in determining T cell fate and (dys)function. Conversely, studying CC in T cells poses other challenges due to a lack of tools for deep CC analysis. CC effects in single-cell RNA-sequencing are often regressed out as a confounding variable^32^, and methods for CC interrogation from transcriptomics are not suited for deep CC molecular analysis because CC checkpoints are intrinsically controlled by protein abundance and post-translational modifications.^33–35^ Fluorescent microscopy uses fluorescent-tagged CDT1 and Geminin to image CC progression^36^, but these methods are limited by low-dimensional and low-to-moderate-throughput assays, and microscopy is not well-suited for asynchronously dividing cell suspension systems such as primary human T cells. Additionally, expression of CDT1, Geminin, and other CC-related molecules can exhibit noncanonical expression patterns^37^, making these low parameter methods unreliable for accurate estimation of CC timing in dysfunctional cells. Fluorescence flow technologies identify proliferating cells and annotate cell cycle phases^38,39^ but CC phase annotations are coarse and do not fully capture the continuous and dynamic nature of cellular transitions as a continuum, which is pivotal for understanding the intricate link between CC timing and cell development in response to perturbations. Thus, deep estimation of accurate CC progression in dysfunctional cell states relies on multivariate measurement of diverse CC molecules.

To capture the resolution needed for understanding the mechanistic link between CC programs and T cell (dys)function, we leveraged single-cell mass cytometry^40^ (scMC) for simultaneous measurement of diverse CC regulators, proliferative tracing, lineage markers, and functional proteins across millions of primary human T cells over successive proliferative generations, multiplexed across time and perturbations. We employed uniquely designed probes that capture both canonical and noncanonical (aberrant) CC states as well as T cell activation, inhibition, and differentiation states. We further bridge the cell-state gaps between multiple static snapshots, over time, using trajectory inference^41^ (TI) to create continuous pseudotimes of cell cycle transitions and T cell fate. This enabled a deep interrogation of the link between CC decisions and T cell (dys)functional states. To help decode this relationship, we introduced a time-series computational framework called Hafez that detects and quantifies trajectories in the presence of aberrant cells and perturbations like kinase inhibition, as well as a post-trajectory inference analysis toolkit for integrated analysis of both CC and T cell fate trajectories over time and division in mechanistic studies of T cell (dys)function.

We modulated CC with cyclin-dependent kinase (CDK) inhibitors, an interleukin-2 inducible T cell kinase blockade, and with a tonic signaling chimeric antigen receptor T cell model of exhaustion to disentangle the crosstalk between CC sensing and T cell differentiation. With this we demonstrate that CC sensing during T cell activation augments T cell differentiation where CC slowing combined with TCR signaling blockade has additive or synergistic consequences on T cell state. We additionally demonstrate that tonic signaling in T cells drives aberrant proliferative CC states that persist in exhausting T cells even when receptor signaling is shut off. These aberrant CC signatures also persisted *in vivo* 413B-engrafted mouse models with human CAR TILs and *in situ* human pan-cancer exhausted CD8 TILs, with key CC phase-dependent gene programs associated with deepening exhaustion signatures. Our findings show that aberrant, proliferative CC arrest programs are intrinsically linked upstream in the determination of T cell (dys)functional fate.

## Results

### Decoding cell cycle dynamics in primary T cells over time and division

CC entry and proliferation is a hallmark result of TCR stimulation. Studies have investigated the tight link between CC and T cell differentiation as a function of time-since-activation or division, implicating proliferation and CC dwell times in T cell fate.^1,12,19,26,27,33,34,42^ Still, much remains unknown about the relationship between CC and T cell state. This is due to experimental and technological gaps in measuring accurate, granular CC progression over multiple timepoints, divisions, and multi-dimensional readouts of CC and T cell state in suspension cells. To study primary human T cells across these variables, we used single-cell MC and implemented Hafez for trajectory inference and post-trajectory analysis (**Figure S1, methods)**. We stimulated CFSE-labeled primary human T cells and created a model of T cell activation, tracing multiple generations of cells over time, division, and CC-phase using mass cytometry. In doing so, we captured the profile of ∼14 million cells across 9 divisions (i.e. generations), spanning 10 days, and 3 healthy blood donors. Based on this matrix of features we had an analytic space of 216 (timepoint, cell-state, Division) conditions for each single cell feature captured (**Figure 1A, S2A-B)**. This enabled us to map the influence of CC timing and division on human t cell fate choices in a manner that has not been previously possible.

**Figure 1:**
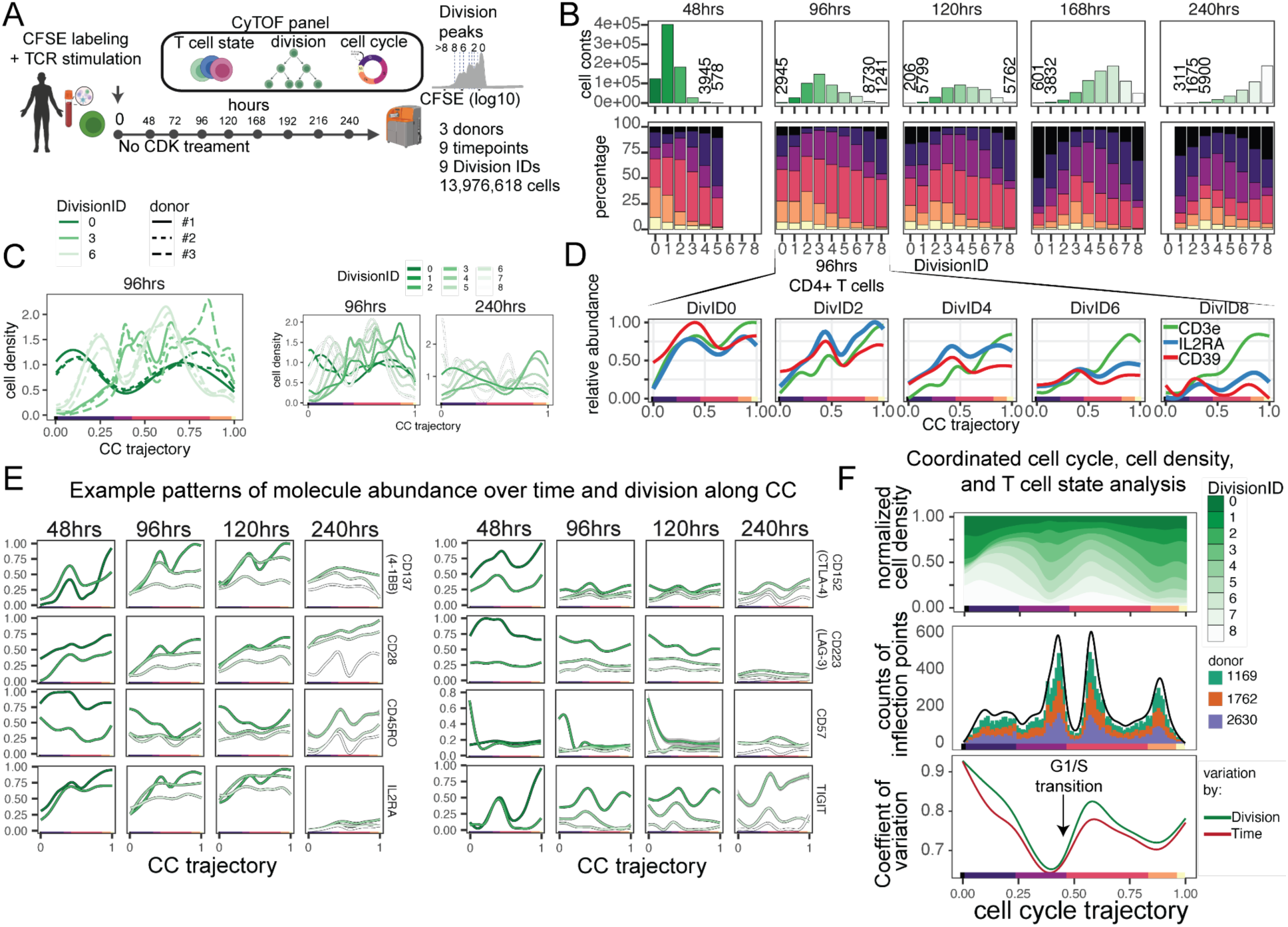
A Mass Cytometry platform to deeply quantify cell cycle progression and division, over time, with primary human T cell states. **(A)** Experimental design for ex vivo T cell stimulation with CFSE Division IDs (n=3 donors) in CD3/CD28 stimulated primary human T cells. **(B)** Cell counts and CC phase fractions over time and division. **(C)** CC occupancy curves across different donors. **(D)** CD3, IL2RA, and CD39 example molecular dynamics for multiple division IDs at 96 h. **(E)** Smoothed oscillation patterns for different T cell molecules along cell cycle pseudotime with time and division. **(F)** Coordinated cell density across divisions, inflection point analysis, and covariance analysis of T cell phenotypic identity at all timepoints and divisions at 96 h. **(G)** dDR^Hafez^ embedding of distributions, each point is one distribution. **(H)** Relative enrichment scores for different T cell targets comparing cells with high expression relative to cells with low expression. **(I)** Network plot (nondirectional) of 30 strongest scores biasing CC occupancy for interacting T cell state molecules. Size of lines indicate strength of the interaction.

### CC phase dynamics in primary T cells are linked to time, division, and T cell molecule expression

We previously showed that early differentiation choice is linked to time and proliferation in *ex vivo* activated T cells using CFSE proliferation tracing, but the role of direct CC phase influence T cell state, remained unknown.^19^ Thus, we first estimate cell distributions along the CC to uncover CC dynamics with division- and time-dependencies. Cell density along CC pseudotime is robustly division-dependent, with average pseudotime clearly staggered with each division **(Figure S2C)**. Deeper analysis reveals coordinated timepoint- and division-dependent CC phases where maximum proliferative potential (non-G0) is achieved between 96h and 120h, with a return to G0-like and G0G1 states at later divisions, and later timepoints post-stimulation (**Figure 1B, S2D-E)**. The cell density patterns along the CC trajectory were remarkably robust for different donors (**Figure 1C)**, reinforcing the idea of CC phase as a conserved regulatory feature.

After the initial proliferative burst, both the remaining slow dividing cells and rapid dividers are biased towards G0/G0G1 CC states–indicative of their slowed division rate–and by day 10 most cells experience >8 divisions and cannot be traced. Notably, earlier divisions enriched towards late S phase and later divisions biased towards early S phase, suggesting differential CC phase transit controlled by division state after stimulation **(Figure S2F)**. Paired with this coordinated CC transit, the molecular abundance of T cell targets also oscillates with time and division, increasing or decreasing at different points along the trajectory (**Figure 1D-E)**. Collectively, these findings reveal that CC transit is an actively remodeled feature of T cell activation, shaped by both elapsed time and division history as cells re-enter into quiescent-like states after generations.

Considering this observation that CC phase dwell time is re-modeled with time and generation, (**Figure 1C, S2D-E)** and that cell cycle transit can control cell fate mechanisms in multipotent cells^1^, T cells^26,42^, and other hematopoietic lineages^43–45^, we sought to specifically investigate the relationship between CC transit with indicators of T cell activation, co-stimulation/co-inhibition, senescence, and differentiation. Thus, we use Hafez to count the number of inflection points (CC-associated changes in expression dynamics) in T cell markers and map this to cell density over time and division **(Figure S2G)**. CC density distributions occupied both highly transient and highly dense CC states along the trajectory, suggestive of differential CC transit dwell-times (**Figure 1F, top)**. By counting inflection points of T cell phenotypic markers along the CC trajectory, we observed CC-associated switches during mid-G0G1 and striking inflections at the G0G1-to-G1 transition, during early S phase, and just before the S/G2 checkpoint (**Figure 1F, *center*)**. Inflections in T cell protein expression were highly coordinated with CC density regions that were either very high (long CC dwell times) or very low (transient CC states) (**Figure 1F, *top and center*)**. Particularly, the G0G1/G1 and S/G2 switch points were a highly transient CC state characterized by lower cell density, while the mid-G0G1 and early S phase switches were preceded by higher cell densities. Finally, T cell, diversity defined by the average coefficient of variation over time or division for T cell phenotypes, coincided with these CC inflection points, suggesting that T cell diversity during differentiation substantially changes with this exact CC-phase timing (**Figure 1D, *bottom*).** Notably, the lowest point of variation occurred just before the rapid G1/S transition, which is a critical checkpoint of cell differentiation in human embryonic stem cells, neural stem cells, and other multipotent cells.^4,19,46,47^

Overall, by incorporating CC-phase granularity with long-term generation tracing and quantitative trajectory modeling, we show that T cell phenotypic diversity is influenced by CC timing, where fewer T cell state changes before, but greater remodeling occurs after, the G1/S transition **(Figure S2H)**. Taken together, our results suggest that T cell differentiation and fate decisions upon stimulation may be intrinsically linked to CC-phase timing where progression directly dictates fate decisions.

### *In silico modeling of* CC timing and T cell state molecules establishes upstream, intrinsic link of CC transit to human T cell fate

To test whether CC phase occupancy is a structured feature of human T cell differentiation rather than a stochastic byproduct of activation, we leveraged our large dataset of primary human T cells spanning multiple donors, timepoints, division states, and discretized cells into high and low expression of T cell targets. We also included cycling dependent kinase (CDK) inhibitor treatments as a positive control for stimulated but arrested T cells. Using dDR embedding of CC-phase distributions for each group, we compressed approximately 15 million cells into 14,000 points in a low-dimensional landscape where proximity reflects similarity in CC occupancy patterns across the combination of all these groups — effectively capturing the CC “state space” of human T cell activation at scale (**Figure 2A)**. Critically, clustering within this space was not random: arrested states from unstimulated and CDK-inhibitor-treated cells co-localized, as did highly proliferative subsets from early stimulation timepoints, G0/G1 checkpoint accumulation, and distinct S and G2 phase buildups — confirming that CC-phase distributions faithfully reflect known differences in T cell activation state rather than technical or donor-driven variation (**Figure 2B)**.

**Figure 2:**
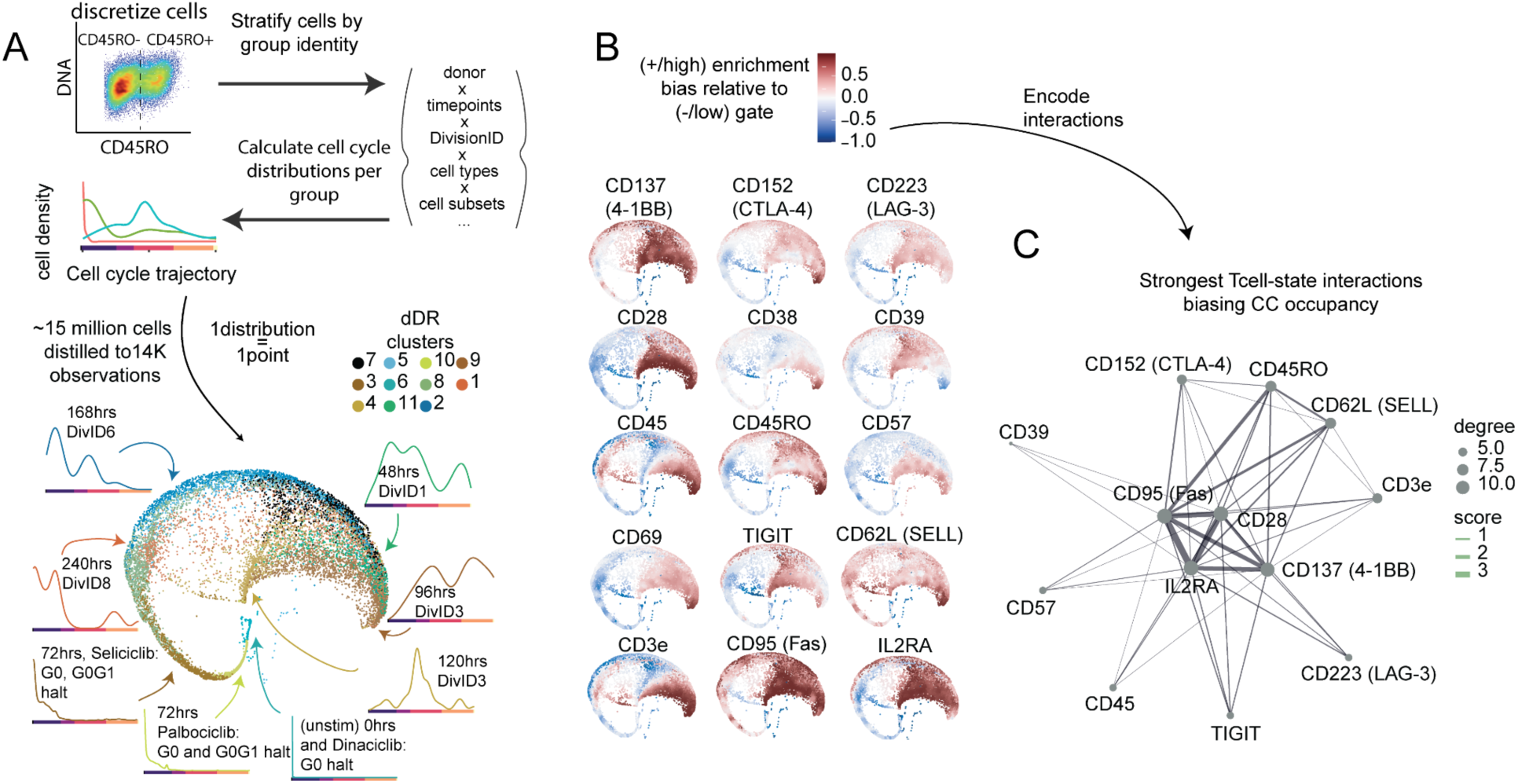
CC phase occupancy in activated human T cells is structured by T cell differentiation state. **(A)** dDR^Hafez^ embedding of distributions of T cell-cell cycle trajectories (n=3 donors). Here, each datapoint represents a CC density distribution for one of the approximately 14,000 conditions. Groups of cells used to compute the CC density distribution are stratified by donor, timepoint, division, CD4/CD8 identity, and discretized T cell state markers. Conditions with similar CC distributions are closer in the embedding space. A cell cycle arrest experiment with CDK inhibitors (n=3 donors per treatment) is included in the embedding analysis as a positive control for non-proliferative groups in stimulated T cells (Palbociclib, Dinaciclib, Seliclib). **(B)** Relative enrichment scores for different T cell targets comparing cells with high expression relative to cells with low expression. **(C)** Network plot (nondirectional) of 30 strongest scores biasing CC occupancy for interacting T cell state molecules. Size of lines indicate strength of the interaction.

Having established that CC occupancy is structured, we next asked which T cell molecules most strongly bias it. Activation and costimulatory signals — CD25, CD28, CD137, and CD95 — emerged as the dominant and most promiscuous drivers, biasing cells toward distinct proliferative CC states across diverse molecular co-expression contexts (**Figure 2C)**. Memory and early differentiation markers CD45RO and CD62L showed weaker but similarly broad influence. Strikingly, inhibitory molecules including CD57, CTLA-4, and LAG-3 interacted far more selectively — and nearly always converged on CD95/FAS co-expression — directly linking CC state bias to the apoptotic and exhaustion machinery governing T cell dysfunction (**Figure 2C)**. Together, these data establish that CC phase occupancy in human T cells is a structured, cell-intrinsic feature of the differentiation program, shaped most strongly by activation state and constrained by inhibitory signaling through apoptotic checkpoints — nominating CC transit itself as a candidate regulator of T cell fate.

### Transient CC inhibition during T cell activation durably reprograms differentiation trajectory

With our observation of the influence of CC entry, dwell time and coordination around the G1/S on T cell state (**Figure 1-2**), we hypothesized that earlier, targeted CC-phase slowing could causally influence T cell differentiation. To dissect this and probe the mechanistic influence of CC on T cell state, we isolated primary human naive T cells and screened various CDK inhibitors (CDKIs) which slow CC-phase progression during anti-CD3/anti-CD28 stimulation. On top of our CC analysis framework, we tracked 22 T cell molecules spanning activation, co-stimulation, inhibition/senescence, collected T cell subsets on days 3, 7, 15, and 22 post-stimulation, discretized six CFSE-based cell divisions, and computed parallel CC and CD8 differentiation trajectories capturing the naive-to-Temra continuum (**Figure 3A-B, S3A-D).**

**Figure 3:**
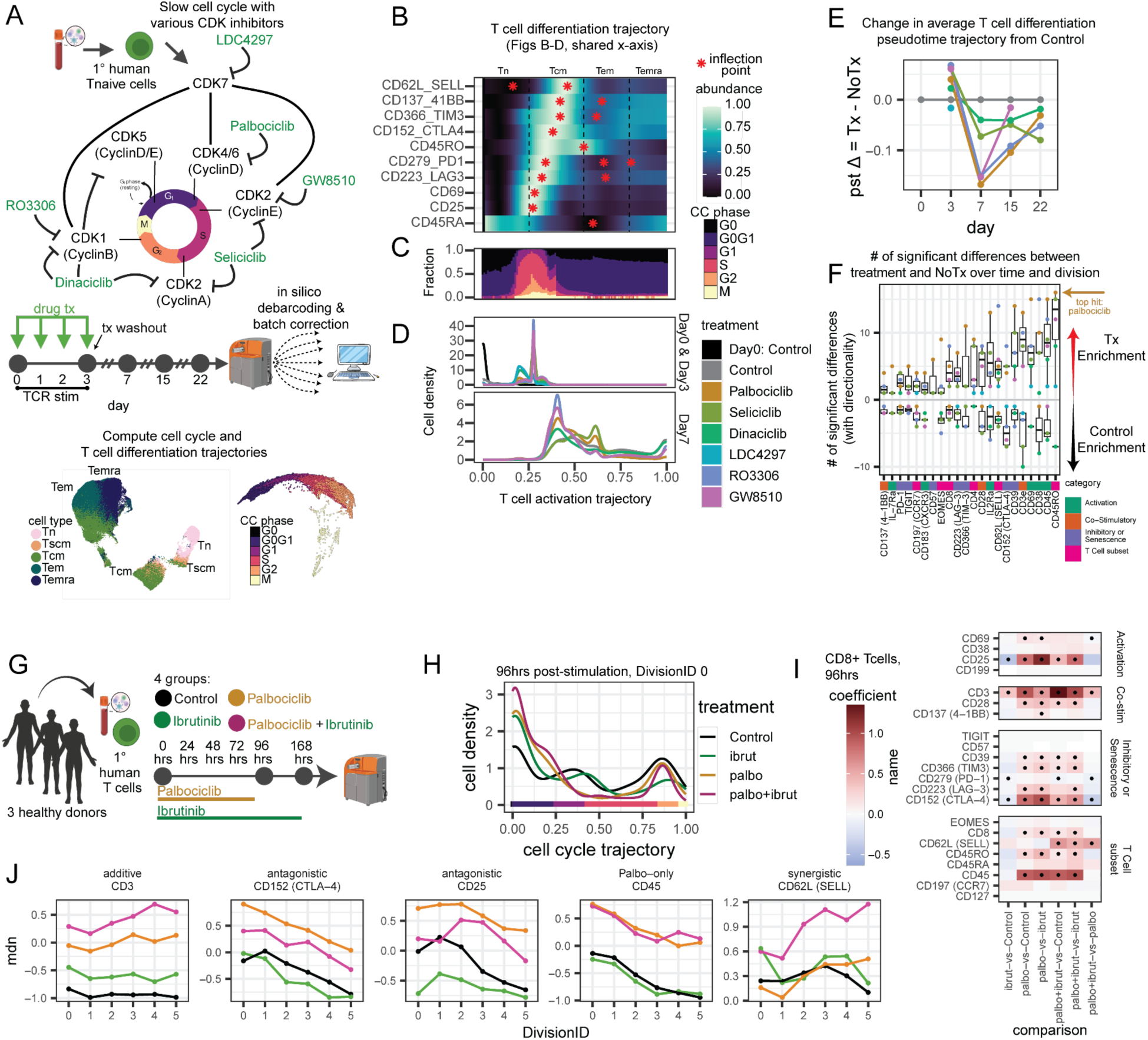
Earlier CC sensing and CD3/CD28 co-stimulation crosstalk has a durable effect on modulating T cell fate. **(A)** Diagram of CDK inhibitor targets in T cells, and experimental design for ex vivo TCR stimulation screening CDK inhibitors on days 0, 3, 7, 15, and 22 (n=2 donors). Embedding of cell cycle features colored by CC phase and embedding of T cell features colored by cell type or T naive (Tn), T stem cell memory-like (Tscm), T central memory (Tcm), T effector memory (Tem), and T effector-memory re-expressing CD45RA (Temra). **(B-D)** T cell differentiation trajectory on x-axis shared across figures C-E: **(B)** Heatmap of smoothed expression for each target related to T cell differentiation along T cell differentiation trajectory. Red points indicate an inflection point. **(C)** Cell cycle phases along CD8 T cell differentiation trajectory at D3. **(D)** Cell density^Hafez^ along T cell differentiation trajectory at day 0, 3, and 7 for all conditions. **(E)** Change in average T cell differentiation trajectory across all cells for each treatment compared to Control for each day. **(F)** # of significant differences (padj<=0.05) comparing each treatment condition compared matching across time and division (see methods). Positive values indicate increased molecular abundance in the specified treatment compared to Control and negative values indicate decreased molecular abundance compared to Control. **(G)** Diagram of experimental design for ex vivo T cell stimulation with CDK4/6 inhibition using palbociclib, ITK blockade using Ibrutinib, and combination. **(H)** Cell cycle density of undivided cells 96hrs after bead stimulation. **(I)** Differential abundance analysis of T cell features comparing treatments for CD8^+^ T cells at 96hrs after bead stimulation matched for donor and DivisionID with dots indicating significance (adjusted p-value <= 0.10). **(J)** Median value for example significant T cell features from (K) over divisions at 96hrs.

As expected, CC inhibition enriched for T cell states immediately before G1 entry and early Tcm state along the differentiation trajectory on day 3 (**Figure 3C-D).** This early Tcm enrichment was tightly associated with proliferative CC signatures, increase of activation-(CD69, CD25) and inhibition-related molecules (PD-1, LAG-3, CTLA-4) along the T cell differentiation trajectory. Notably, undivided cells experienced a rapid G1-dependent remodeling for many T cell markers, including modulation of expression for CD25, memory (CD45RO, CD62L), and early activation markers (CD45, CD69) **(Figure S3E).**

Despite all CDKi treatments washed out after D3, differentiation skewing persisted robustly through days 15 and 22 — demonstrating that a transient perturbation of CC transit during the activation window is sufficient to durably reprogram T cell fate (**Figure 3E)**. This lingering effect was most clearly reflected in memory marker remodeling, with CD45RO, CD62L, and CD69 among the most consistently altered molecules across inhibitors, timepoints, and Divisions (**Figure 3E, S3E).** Across CDKi treatments, the dominant fate outcome was enhanced memory formation: palbociclib, seliciclib, and ro3306 skewed cells toward early effector memory (Tem)-like states, while dinaciclib favored late Tcm states — but all converged on a net increase in CD45RO+CD45RA- cells averaging 30.7% in CD4 and 20.5% in CD8 T cells by day 22 30.7% in CD4 T cells and 20.5% in CD8 T cells (CD45RO+CD45RA- % increase in CD4: palbociclib 32.0%; seliciclib 36.9%; dinaciclib 13.3%; ro3306 40.7%. CD8: palbociclib 28.7%; seliciclib 43.9%; dinaciclib 31.7%; ro3306 47.6%) **(Figure S3H)**. Most inhibitors, despite different CDK target profiles, produced a similar memory-biased outcome that suggests this is a general consequence of CC slowing rather than an on-target effect of any single inhibitor.

Across the inhibitor panel, the strength of fate remodeling tracked with the CC stage being targeted: palbociclib, which selectively inhibits CDK4/6 and arrests cells at the G0/G1 and G1/S boundary, was consistently the strongest modulator of T cell differentiation state (**Figure 3F, S3F-G**). In contrast, inhibitors acting on downstream CC kinases — later in the cycle — produced weaker and less consistent fate skewing. This hierarchical pattern argues that the G1/S restriction point is not merely one of several permissive windows, but the primary checkpoint at which CC transit speed is translated into a differentiation decision.

Together, these data support a model in which CC transit speed during the TCR activation window acts as a direct rheostat for T cell fate: slowing progression transiently enhances memory formation at the expense of terminal effector differentiation, and this fate bias is set early and maintained. Given that this window coincides with the G1/S restriction point identified in our correlative analysis (**Figure 1–2**), CC transit at this checkpoint emerges as a candidate causal mechanism–rather than a passive correlate–of T cell differentiation decisions.

### Cell cycle inhibition and TCR signaling blockade act synergistically on T cell programs

Considering that CC inhibition slows T cell differentiation and early TCR signaling blockade via ITK inhibition slows CC, we sought to understand whether the effects of ITK inhibition on T cell state may also be explained by CC slowing. To assess this, we treated CFSE-labeled *ex vivo* expanded total T cells with ibrutinib (ITK blockade), palbociclib (CDK4/6 inhibition), or both (**Figure 3G)**. All treatments slowed without halting CC progression during TCR activation **(Figure S3I-J)**. Ibrutinib and palbociclib caused similar shifts in CC distributions in undivided cells, and combination treatment caused even greater G0G1 skewing (**Figure 3H)**.

CC inhibition and TCR signaling blockade reshaped CD8 T cell states through additive, antagonistic, and synergistic interactions. While individual drugs alter activation, costimulation, inhibitory, and subset markers, their combination produces non-linear outcomes. For example, CD3 downregulation was additively suppressed by both inhibitors, CTLA-4 and CD25 showed antagonism, CD62L increased with combined treatment, and CD45 loss was mitigated only by palbociclib – which is consistent with ITK blockade being downstream of CD45 in TCR signaling. Thus, dual blockade does not simply add effects but rewires T cell regulation in drug-specific ways, and the effects of TCR signaling blockade is partially enhanced by cell cycle inhibition.

Taken together, these results underscore the crosstalk between CC sensing and TCR signaling. While some of the effects of CC sensing and TCR signaling blockade are partially explained by each other, our data provides evidence that T cells integrate signals in tandem which has direct consequences on the resulting cell state progression. Thus, CC sensing during the stimulation phase can lead to cell state biases that are imprinted even in the presence of TCR signaling blockade, highlighting the intrinsic link between CC and T cell differentiation.

### Tonic signaling drives aberrant S/G2 and G2/M accumulation in T cell exhaustion

Based on inhibitory/senescent T cell molecules showing bias to specific CC phase patterns (**Figure 1–3**) and the role of dysfunctional TCR signaling associated with T cell exhaustion, in general, we investigated whether there could be a CC phase component of this T cell behavior. In primary T cells, expression of inhibitory receptors and the overall exhausted T cell state can result from persistent antigen exposure, resulting in loss of effector functions and proliferative capacity.^13^ To consider this connection between persistent signaling, CC aberrancy, and emergence of T cell exhaustion features, we used our CC platform with Hafez time-series analysis to quantify the relationship between tonic signaling, exhaustion, and the CC state. To that end, we use a chimeric antigen receptor (CAR) HA-28z tonic signaling model that drives T cell exhaustion versus an equivalent CD19 CAR-T as a model system^48^. Upon activation and transduction (**Figure 4A)**, HA CAR-T cells exhibited a robust accumulation of the mitotic-related S/G2 and G2/M phases compared to CD19 and MOCK transduced at day 10, despite the reduced proliferative capacity of HA CAR-T cells (**Figure 4B)**. Additionally, HA CAR-T cells exhibited aberrant CC phenotypes in early (G0, G0/G1) as well as late (S/G2, G2/M) CC timing, with aberrant expression of the mitosis-initiating proteins CyclinB1, pRb, and PLK1 (**Figure 4C-D)**. Notably, PLK1, crucial for cell division^49,50^, is atypically high in G0, G0G1. HA CAR-T cells also had a global decrease in Geminin expression, a key regulator of DNA replication before S phase initiation.^51^ Thus, this S/G2 and G2/M accumulation, revealed by Hafez robustly across eight donors (**Figure 4E)**, establishes that dysfunction tonic cell signaling can directly drive abnormal CC phase ocupancy in a consistent way in human T cells.

**Figure 4:**
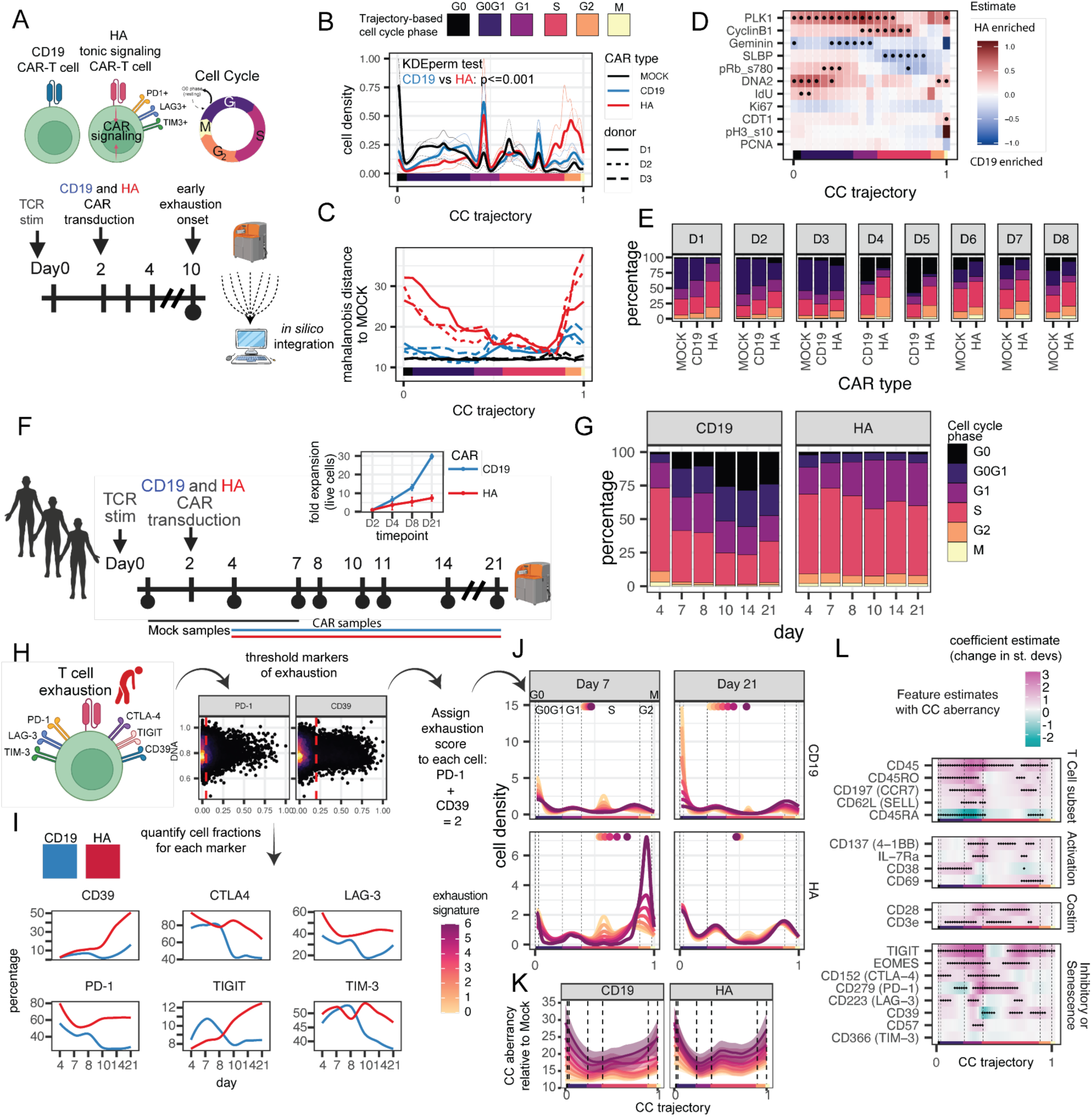
Tonic signaling biases cells towards aberrant, proliferative CC signatures. **(A)** Cartoon schematic of CD19 and HA CAR T cell transduction experiment for scMC analysis. **(B)** CC trajectory for MOCK, CD19, and HA CAR T cell types across 3 donors on day 10 after anti-CD3/anti-CD28 stimulation. KDE permutation test CD19 vs HA, p<=0.001. **(C)** Cell aberrancy score for CD19 and HA CAR T cells compared to MOCK landmarks. **(D)** Significant trajectory-matched CC features between HA and CD19 CAR T cells. Point indicates padj<=0.1. **(E)** Deep CC phase annotation across 8 donors for MOCK and/or CD19, and HA. **(F)** Experimental design to track CC signatures over time **(G)** CC phase annotations over time for CD19 and HA CAR T cells. **(H)** Cartoon schematic and gating scheme of features of exhaustion and exhaustion score. **(I)** Fraction of positive cells for each exhaustion feature. We assigned an exhaustion score to each cell depending on their marker positivity by computing an aggregate score of 0-6, and then in silico extract cells from each aggregate score to study their CC distributions. **(J)** Cell density along CC trajectory for CD19 and HA cells from each exhaustion score. **(K)** Smoothed gam of CC aberrancy stratified and colored by aggregate exhaustion count scores. KDE linearity score between CC aberrancy and ordinal exhaustion scores p<=2e-16 (see methods). **(L)** Estimated aggregate coefficient change associated with cc aberrancy along the CC trajectory for each feature across all timepoints, donors, and CAR types. Black dots indicate statistical significance (see methods) for padj<=0.1.

### Tonic signaling sustains an exhaustion-associated CC aberrancy over time

To better understand the long-term consequences of tonic signaling associated with CC aberrancy on the onset of exhaustion, we followed CD19 and HA CAR-T cells until day 21 of expansion (**Figure 4F)**. CD19 CAR-T cells acquired CC exit signatures at day 10 through day 21, but HA CAR-T cells retained an aberrant, proliferative CC signature despite their poor *in vitro* expansion (**Figure 4F-G)**.

While tonic signaling in HA CAR-T cells is known to drive features of exhaustion, we asked whether these exhaustion signatures are directly associated with the emergence of differential CC states. To do this, we assigned thresholds to quantify a score using markers of T cell exhaustion (CD39, CTLA-4, LAG-3, PD-1, TIGIT, and TIM-3) (**Figure 4H)**. Here, we observed an increase in exhaustion of HA CAR-T cells compared to CD19 CAR-T cells at later timepoints (**Figure 4I)**. On D7, before the onset of exhaustion, cells with a higher exhaustion score had more proliferative CC signatures with an S and G2 skew, whereas cells with a lower score had biases towards early S phase or CC exit (G0 and G0G1) (**Figure 4J**). Notably, CD19 CAR-T cells with low exhaustion signatures enriched towards non-proliferative G0 and G0G1 CC signatures at later timepoints and CD19 CAR CC distribution with higher exhaustion scores resembled tonic signaling HA CAR.

Notably, irrespective of CD19 or HA CAR, higher exhaustion scores were also associated with greater CC aberrancies (**Figure 4K**), further enforcing the entrenchment of these phenomena within the biology of the T cell itself. Moreover, these cells with greater CC aberrancies had significantly altered CC-dependent expression of diverse costimulatory, inhibitory/senescence, activation, and T cell differentiation markers (**Figure 4L)**. At the same time, inhibitory/senescence markers like TIGIT, CTLA-4, PD-1, and others had increased expression with CC aberrancy, while differentiation markers like CD45RA had decreased expression particularly restricted to G0 and G1 phases. These T cell molecular shifts with independently calculated CC aberrancy correlates T cell programs with CC phase. Altogether, this tonic signaling exhaustion model demonstrates the direct connection in the aberrancy in specific CC phases that precedes the transition to a dysfunctional T cell fate.

### Signaling rest restores CC phase dynamics in tonic signaling T cells

Considering that the CC aberrancy observed in the HA CAR-T model is sustained over time with tonic signaling, we hypothesized that a reversal of tonic signaling, which can rescue exhaustion, could also restore normative CC phase transitions. Dasatinib–a tyrosine kinase inhibitor that inhibits both TCR and CAR signaling^21,52,53^ can reverse functional and phenotypic hallmarks of exhaustion in this model. To address this, we shut off receptor signaling in tonic signaling HA CAR T cells starting on either D4 or D7 of tonic signaling and track both cell cycle pseudotime and exhaustion molecules for multiple timepoints up to D21 post-stimulation in 3 human donors (**Figure 5A).**

**Figure 5:**
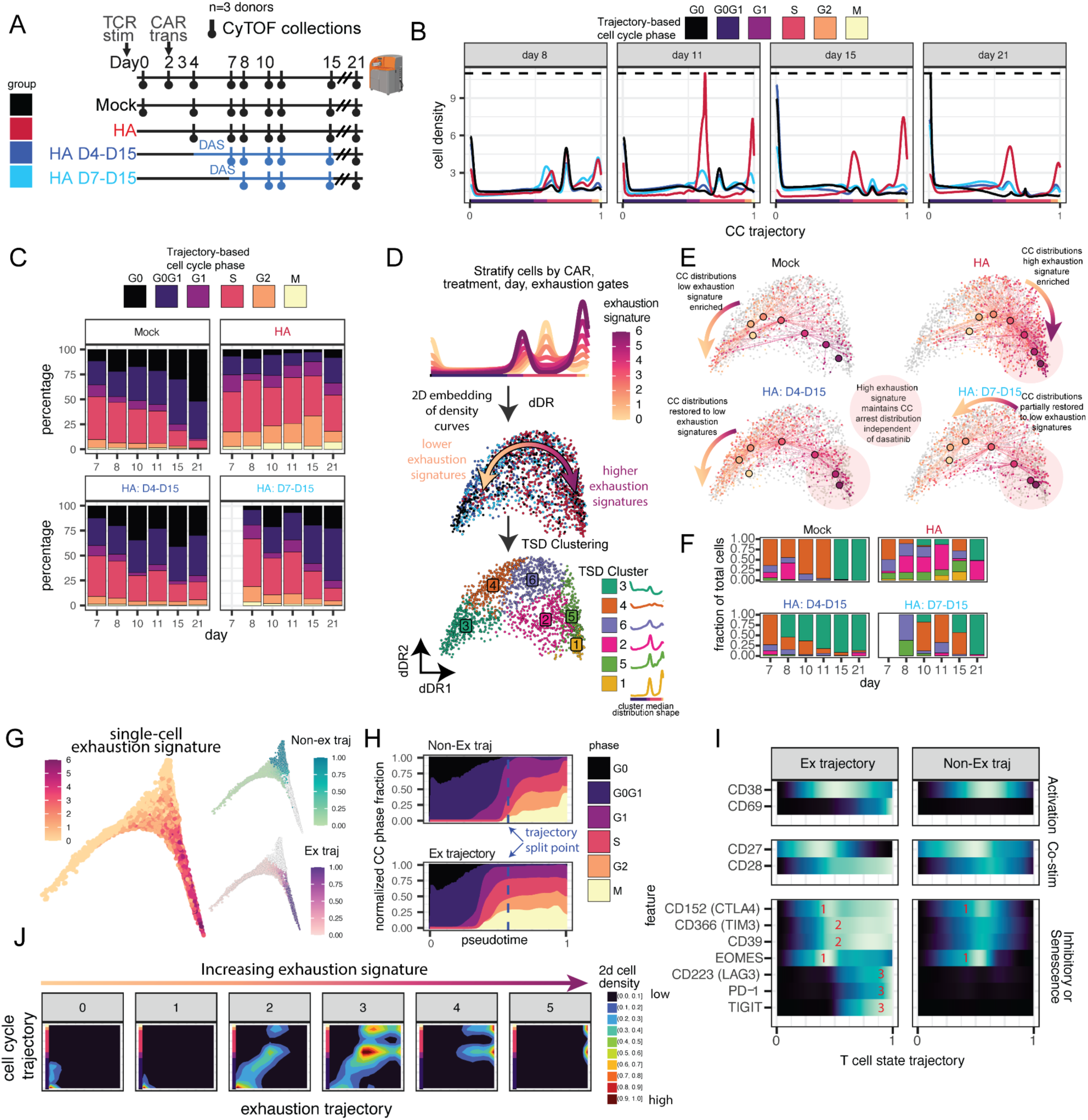
Transient signaling rest mends aberrant CC behaviors and reveals unique signature linked to exhaustion. **(A)** Cartoon schematic of transient rest experiment using Dasatinib in tonic signaling HA CAR-T cells (n=3 donors). **(B)** Mean cell density along CC trajectory^Hafez^ for MOCK, HA, and dasatinib-treated HA D4-D15, and D7-D15 CAR-T cells across 3 donors over time. **(C)** CC phase annotations over time. **(D)** Cartoon of (top) cell density curve along CC trajectory stratified by CAR type, treatment, day, and exhaustion score combinations, (center) dDR^Hafez^ embedding colored by CAR treatment group from (A), and (bottom) TSD clustering results mapped onto dDR^Hafez^ embedding. Each point is a CC density distribution from a group of cells. **(E)** TSD embeddings for each condition colored by exhaustion score. **(F)** Fraction of cells occupying a TSD cluster for each timepoint and group. **(G)** Diffusion map embedding of CAR-T cells colored by exhaustion signature (left). Non-exhaustion (top) and exhaustion trajectories (bottom) are colored in by pseudotime. **(H)** Cell cycle phase fractions along exhausted and non-exhaustion trajectories. **(I)** Molecular dynamics along exhaustion and non-exhaustion trajectories, with numbers indicating order of abundance increase along trajectory. Numbers specific approximate point of peak molecule abundance along the trajectory. **(J)** Cell cycle versus exhaustion trajectory faceted by increasing exhaustion signatures and colored with a 2D kernel density.

Tonic signaling CAR T cells showed early signs of CC aberrancy on D8 and persistent aberrancy through D21, while inhibiting tonic signaling with dasatinib on D4 showed early signs of reversed CC cell density patterns on D8 and later that match mock T cells (**Figure 5B-C, S4A-B).** Starting dasatinib treatment on day 7 showed early signs of restoration of CC cell density and decreasing exhaustion signature on day 10. Shutting off receptor signaling also reduced average CC aberrancy, though CC aberrancy continued to increase in cells with higher exhaustion signatures **(Figure S4C-D).** Thus, shutting off receptor signaling that drives exhaustion emergence also restored CC-phase distributions in tonic signaling HA CAR T cells.

### Early CC entry and G2/M buildup are hallmarks of T cell exhaustion independent of TCR

While dasatinib treatment restored normative CC phases globally, a subset of cells retained high exhaustion signatures. We thus leveraged these high exhaustion signature cells to examine the CC consequences of this dysfunctional state even when tonic signaling is inhibited. As before (**Figure 4H-I)**, we performed *in silico* gating of molecules associated with exhaustion. We generated exhaustion scores for all related molecule combinations and stratified cells into highly multiplexed groups based on: exhaustion score (64 combinations), days (6), CAR types (2), treatments (3), donors (3), and car status (2) to compute a combinatorial space of 4,608 unique CC density distributions. We then used the Hafez framework to perform time-series analysis by squeezing the highly multiplexed CC density distributions into dDR*^Hafez^* for visualization and TSD clustering*^Hafez^* to discover patterns in CC density distributions (**Figure 5D)**. The dDR*^Hafez^* embedding organized cell density distributions along a continuum, separating low- and high-exhaustion signatures along with treatment conditions, and identified 6 TSD clusters (**Figure 5D, S5A-C)**. Interestingly, the embedded data points representing each distribution from these highly multiplexed groups, regardless of dasatinib treatment, were organized by a spectrum of low to high exhaustion signatures (**Figure 5D-E).** The distributions for higher exhaustion scores maintained late CC distribution biases and aberrancies, which were enriched in the HA untreated cells despite their lower proliferative potential (**Figure 5E, Figure S5C-D)**. Starting dasatinib on D4 shifted most cells into less exhausted signatures (TSD cluster3), whereas D7 treatment had mixed CC restoration outcomes (TSD cluster 4,6,5) (**Figure 5F)**. TSD cluster1 had the greatest CC buildup signature, was enriched for HA untreated conditions, and had the highest average enrichment signature for all exhaustion markers, particularly CTLA-4 **(Figure S5E)**. Notably, progression along the dDR embedding through the TSD clusters correlated with increasing enrichment patterns for different exhaustion molecules, which may suggest different exhaustion molecules arise earlier or later as cells progress through the exhaustion program **(Figure S5E).** Here, cells with increasing exhaustion scores retained their CC density signatures even when receptor signaling is shut off with dasatinib as well as mock-transduced T cells, decoupling tonic signaling and exhaustion. Taken together, these data suggest that CC dysregulation is an intrinsic feature of the exhausted state, maintained independently of receptor signaling–but whether exhaustion follows an ordered, predictable program linked to specific CC transitions remains an open question.

To address this directly, we performed unsupervised trajectory analysis of T cell molecules and identified a branching trajectory splitting into exhausted and non-exhausted lineages from a shared origin (**Figure 5G).** Along the shared portion of the trajectory, CC patterning was distinct, with cells destined for the exhaustion lineage showing greater proliferative CC signatures earlier than the non-exhaustion aligned cells (**Figure 5H**). As cells committed to the exhaustion branch, this early proliferative burst was preceded by elevated CTLA-4 and EOMES expression, followed by acquisition of TIM-3 and CD39, and subsequently LAG-3, PD-1, and TIGIT — revealing an ordered molecular program accompanying CC dysregulation (**Figure 5I).** Strikingly, as cells progressed through this exhaustion trajectory, CC restriction at both G1/S and G2/M increased in step with their exhaustion burden (**Figure 5J)**. Overall, these data mechanistically demonstrate a path for tonic or chronic signaling in T cells to lead to exhaustion, where an earlier and aberrant CC entry results initiates an ordered, self-reinforcing program in which progressive acquisition of exhaustion molecules is coupled to increasing CC phase restriction–implicating CC transit not as a downstream consequence of exhaustion, but as an active participant in its progression–implicating CC transit not as a downstream consequence of exhaustion, but as an active participant in its progression.

### Cell cycle biases are linked with T-cell exhaustion signatures in vivo

The observation that CC biases and aberrancies are associated with exhaustion even, in the absence of TCR signals, underscores the tight link between these states. As such, we hypothesized that this relationship could go beyond *ex vivo* expansion where T cell exhaustion signatures (CARTex) could persist with CC abnormalities *in vivo*. To that end, we analyzed scRNAseq of harvested CAR TILs from 143B-engrafted mice^54^. These include tonic signaling (const) CAR TILs and SNIP (snp) CAR TILs, which have regulated CAR activity using an FDA-approved protease inhibitor that shuts off tonic signaling (**Figure 6A)**. While const CAR TILs had increased proliferative cell cycle phases (S, G2M) compared to snp CAR TILs (**Figure 6B)**, which is consistent with our dasatinib HA CAR-T model, it remains unclear whether there are CC phase-dependent genes that drive transcriptional activity towards exhaustion programs.

**Figure 6:**
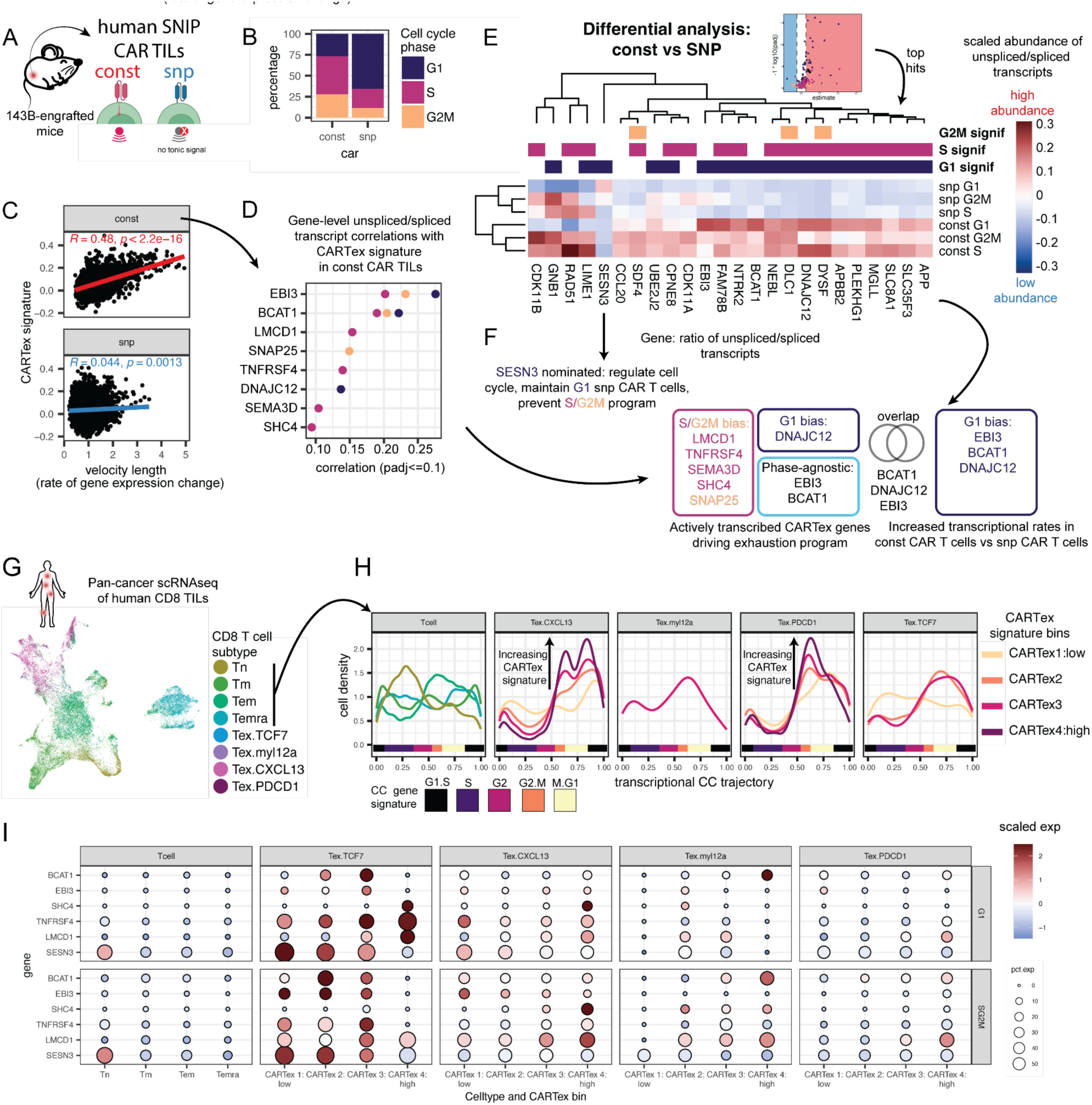
Cell cycle biased exhaustion persists in vivo tumor T cells. **(A)** Cartoon schematic of const and snp CAR TILs harvested from 143B-engrafted NSG mice on day 15 after tumor implantation for single-cell RNAseq. **(B)** Cell cycle phase fractions for const and snp CAR TILs. **(C)** Correlation analysis of CARTex signature and velocity length as a proxy for rate of gene expression change. **(D)** Spearman correlations of CARTex signature with unspliced/spliced transcript ratios for CARTEx genes. **(E)** Heatmap of differential analysis results of unspliced/spliced transcript ratio comparing const vs snp CAR TILs in a phase-matched analysis. **(F)** Nominated gene hits for analysis in (D) and (E), and their overlap colored by CC associations. **(G)** UMAP embedding of CD8 TILs from pan-cancer scRNAseq dataset^53^, including Tn (4,692 cells), Tm (14,546 cells), Tem (13,553 cells), Temra (8,162 cells), and Tex (10,608 cells). Tex cells are further split into Tex.CXCL13 (7,750 cells), Tex.myl12a (246 cells), Tex.TCF7 (504 cells), and Tex.PDCD1 (2,108 cells). **(H)** Transcriptional cell cycle trajectory for different T cell subsets, and exhausted T cell (Tex) subtypes with published CAR Tex signature binned from low to high (1 to 4).

To address whether active transcriptional activity towards exhaustion programs were coupled to CC in const and snp CAR T cells, we correlated CARTex gene signatures **(see methods and Supplementary Table 7)** with active transcriptional reprogramming by leveraging mRNA splice states^5^ to quantify unspliced and spliced transcripts as a proxy for ongoing transcriptional activity and active cell differentiation. Velocity length, a proxy for the magnitude of transcriptional change that reflects the speed of cell state transitions and active gene expression dynamics as inferred from unspliced/spliced transcript ratios, was correlated with greater CARTex signatures in const but not snp CAR TILs (**Figure 6C)**. CARTex genes driving the exhaustion program in const CAR TILs primarily included genes significantly correlated in S/G2 cells; LMCD1, TNFRSF4, SEMA3D, SHC4, and SNAP25 but also included G1-biased DNAJC12 and CC phase-agnostic EBI3 and BCAT1–both of which are implicated in T cell exhaustion (**Figure 6D)**.^55,56^ These directional associations nominate potential drivers of cell cycle-dictated exhaustion programs, however, it remains unclear whether gene activity is driving or compensating for exhaustion here.

To answer this, we compared unspliced/spliced transcript ratios of const versus snp CAR TILs. Const CAR TILs had globally higher gene activity across all phases with signatures of active T cell signaling, sustained proliferation, and cell stress (**Figure 6E)**. This includes increased T cell signaling adaptors (LIME1), metabolic regulators (BCAT1, MGLL, DNAJC12), and cell cycle molecules (CDK11A/B, DLC1) as well as nominated genes previously uncharacterized in exhausted T cells. An interesting exception to the globally increased active transcriptional expression of const CAR TILs is SESN3, which had an increased unspliced/spliced ratio in non-proliferative/G1 snp CAR TILs and is involved in regulating mTOR pathway and providing metabolic stability. This nominates SESN3 as a candidate brake that facilitates and maintains return to quiescence for T cells. In summary, these data point to a layered model of CC-coupled exhaustion in const CAR TILs. Metabolic and stress regulators including BCAT1, EBI3, and DNAJC12 are actively transcribed across all CC phases, while loss of the quiescence-stabilizing brake SESN3 may facilitate sustained CC entry. As cells progress and maintain the CC program, the exhaustion program is amplified–implying that CC phase progression does not merely accompany exhaustion, but actively drives its deepening (**Figure 6F)**.

Finally, we quantified whether these proliferative signatures with exhaustion biases persist in humans T cells *in vivo* by analyzing scRNAseq from CD8 tumor infiltrating lymphocytes (TILs) across 8 human cancers, 45 patients, and 51,561 cells annotated as Tn, Tm, Tem, Temra, and Tex.^57^ To query this more comprehensively, the Tex signatures here are further stratified into 4 subsets (i.e., Tex.myl12a, Tex.PDCD1, Tex.TCF7, and Tex.CXCL13, **Figure 6G)** using the study’s published annotations. We calculated a CAR-Tex score and binned cells with increasing exhaustion scores (1-low, 4-high) **(Figure S5F)**. A transcriptional CC trajectory analysis using our previously published approach captured G0G1-like Tn cells and more proliferative CC states in differentiated T cell subsets. Notably, Tex subsets had a specific CC cell density pattern positively associated with the CARTex signature, like the tonic signaling HA CAR T cell model with a late SG2 buildup (**Figure 6H)**. The exception for this CC bias was the Tex.TCF7 cells with a low CARTex signature (CARTex 1), which do not exhibit this CC state buildup and have more homogeneously distributed CC states. Still, upon increased CARTex signature, the Tex.TCF7 population acquires a CC buildup bias. It has been hypothesized that progenitor cells expressing TCF7 give rise to terminal Tex cells^8,10,57–59^, staging a link for the acquisition of exhaustion between T cell state, exhaustion signatures, and CC fate. Our *in situ SNP CAR-T cell* nominated genes as potential regulators of this linked CC-Ex program also exhibit loss of SESN3 and increased expression of LMCD1 in cells with increasing CARTex signatures and progression through later exhaustion fates after the progenitor Tex.TCF7 state (**Figure 6I)**. This supports a link between exhaustion and aberrant CC programs that work in tandem to drive an exhaustion fate. Taken together, these *in vivo* observations reinforce that alterations to CC transit, CC control, and CC-dependent expression may be a potential precursor to terminal exhaustion fate. Molecular control of CC transit might not simply be a consequence of T cell signaling but directly involved in the trajectory towards T cell exhaustion.

## Discussion

Understanding cell fate choices in healthy or diseased systems requires organizing and deconvolving cell states along time, division, and even across CC within each division. TI has emerged as a crucial tool in this endeavor, allowing us to organize asynchronously behaving cell systems to model cellular transitions across phenotypic landscapes in a quantitative manner to appreciate normative and dysplastic behavior.

In this study, we addressed many of the challenges in multivariate CC modeling during perturbation and introduced useful tools for TI analysis including pseudotime remapping (i.e., DBPN) and quantitative analysis across complex experiments with a multitude of conditions and individuals (**Figure 1-2, S1**). By integrating Hafez with our experimental perturbations of human T cell expansion and multiplexed readout on millions of cells, we have unraveled the intimate link between T cell differentiation, division, and CC dynamics. We demonstrate that this linkage is a generalizable phenomenon existing in the context of CC perturbation, T cell receptor or CAR signaling modulation, or even in unperturbed settings *in vivo*.

We map diverse T cell programs to specific CC patterns that coincide with transient states or dwell times (**Figure 2**). We further demonstrated that this CC sensing-directed cell fate determination during TCR-signaling-based expansion can be modulated by tuning the TCR signaling cascade or CC progression itself (**Figure 3**). Specifically, slowing CC transit with CDK inhibitors during expansion altered T cell output of costimulatory molecules and could enhance less differentiated CD45RO+ memory cells long after activation (i.e., 22 days, **Figure 3**).

In general, CC checkpoints and division are intimately linked to differentiation in pluripotent cell systems, embryonic stem cell differentiation, and hematopoiesis. One overarching theory is that CC phase dwelling can act like checkpoints to serve as phenotypic sinks for T cell program maintenance.^3,22,47,60^ Here, different CC dwell times, induced through varying TCR signaling activation profiles, could be ‘sensed’ through phase-specific gene dosages dictating differentiation decisions. For example, lower affinity receptor engagement with lower magnitude TCR signaling could result in slower CC entry and longer G1/S phase. This type of activation one might expect with a naïve T cell antigen recognition event and our data suggests this type of CC entry and transit would result in a more potent, long lived memory cell formation. On the other hand, chronic activation (i.e., tonic signaling) high affinity TCR engagement, often associated with long-term infection or high dose antigen exposure, could result in pre-mature CC entry and transit with cells dwelling longer at late S/G2/M phase. This exact behavior here was tied to the emergence of more exhausted T cell phenotypes (**Figure 4**). Consequently, these more transient CC states and inflection points in T cell expansion may represent evolved cellular decision points where cells may be most prone to remodeling programs.

Interestingly, the highly transient, G0G1-to-G1 transition was enriched with T cell program inflections (**Figure 2**), suggesting that earlier CC transit may be a window of intrinsic cellular plasticity for T cell fate decisions, before cell fate is locked in. These very same ‘restriction points’ have been studied in pluripotent cell systems, embryonic stem cell differentiation, and hematopoiesis.^4,19,47^ Therefore, it should not be totally unexpected that CC-based mechanisms for cell fate regulation and plasticity exist in T cells as well. If T cells were undergoing the same regulatory phenomena we would expect palbociclib–a CDK4/6 inhibitor–to have the strongest effect on cell state compared to other CDKIs, which indeed was observed.

This is possibly because palbociclib is a highly selective inhibitor of CDK4/6, which tightly controls the G1/S transition and enriches cells in G0G1, compared to other inhibitors that target later-CC-stage CDKs (**Figure 3**).

The idea of CC phase transitions influencing T cell state output is also probable considering the importance of T cell activation and proliferation for achieving effective genetic modification in genetic engineering practices.^61,62^ Moreover, there is also the reported link, with unknown mechanism, between T cell plasticity and regulation of division or CDKs.^19,20,26,27,42^ Finally, the crosstalk between CC regulation and TCR signaling here (**Figure 3)** is further substantiated by the demonstrated additive, antagonistic, and synergistic changes to T cell programs that occur when modifying TCR signaling blockade with ibrutinib, CDK4/6 inhibition, and combinatorial treatment. Given this, it is possible that CC sensing integrating with control of T cell differentiation choices could already be relevant in the context of CDK inhibition as a chemotherapeutic for cancer.^26^

A prime example is exhausted T cells that have both a dysfunctional proliferative capacity and T cell state. Our data shows that these cells maintain an aberrant molecular CC signature juxtaposed with poor cell expansion that is amplified with the highest exhaustion signatures (**Figure 4**), even in the absence of TCR/CAR signaling (**Figure 5**). In these models, downstream CC consequences of tonic signaling-induced exhaustion may be encoded before exhaustion is functionally acquired around day 10 (i.e., perhaps some epigenetic encoding has already set in).

One question that remains is whether CC aberrancy and control work in parallel with T cell signaling to restrict an exhausted fate program. One possible model we can posit that links exhausted T cell differentiation programs with CC derives from the transcriptional signatures of CARTex exhaustion in cancer TILs associated with CC aberrancies (**Figure 6**). We observed that the abundance of Tex cells with S/G2 signatures increased with greater CARTex exhaustion signatures in the more terminally exhausted cell subsets. Conversely, progenitor-like, Tex TCF7 cells, with lower CARTex signatures, did not exhibit S/G2 buildup compared to CARTex-high counterparts. These cell cycle signatures persist in melanoma TILs, where exhausted cells with highest TCR signaling signatures had higher proliferative signatures.^63^ This reinforces the idea that the S/G2 CC buildup program that is acquired with the encoding of exhaustion states is linked to the transition, as opposed to a consequence of the exhaustion state itself. Again, all of this is consistent with the idea of exhaustion being an evolved process that is a response to dysfunctional over-activation of TCR signals, such as chronic LCMV infection, where exhausting cells maintain a proliferative CC signature in earlier timepoints, then when exhaustion programs are complete at later timepoints, cells return to CC quiescence.^59^

Collectively, we demonstrated that our integrated experimental and computational platforms can be used to deeply parse the links between cellular differentiation, proliferation, and CC state across millions of cells and highly multiplexed perturbations or individuals. Considering the pervasive role of CC biology across diverse cell systems and its evolutionary conserved molecular orchestra, CC regulation continues to give credence that it transcends its role as an engine that drives cellular expansion and instead may be directly sensed and integrated into cellular decisions for cellular differentiation. We suspect that other fundamental cell molecular processes (i.e., metabolism, epigenetics, signaling) may further layer over cell expansion and differentiation processes to complement a more holistic regulatory picture. Accordingly, quantitative packages of computational tools for discovery of relationships in single cell trajectories, like Hafez here, will be able to capture the nuances pivotal to understanding cell decision making in normative and disease processes. New computational technologies to help understand these integrative processes will not only deepen our understanding of cellular behaviors but also provide foundational insights required for advancing strategies to effectively engineer therapeutic cells and interventions.

## Supplemental figures

**Supplementary Figure 1.**
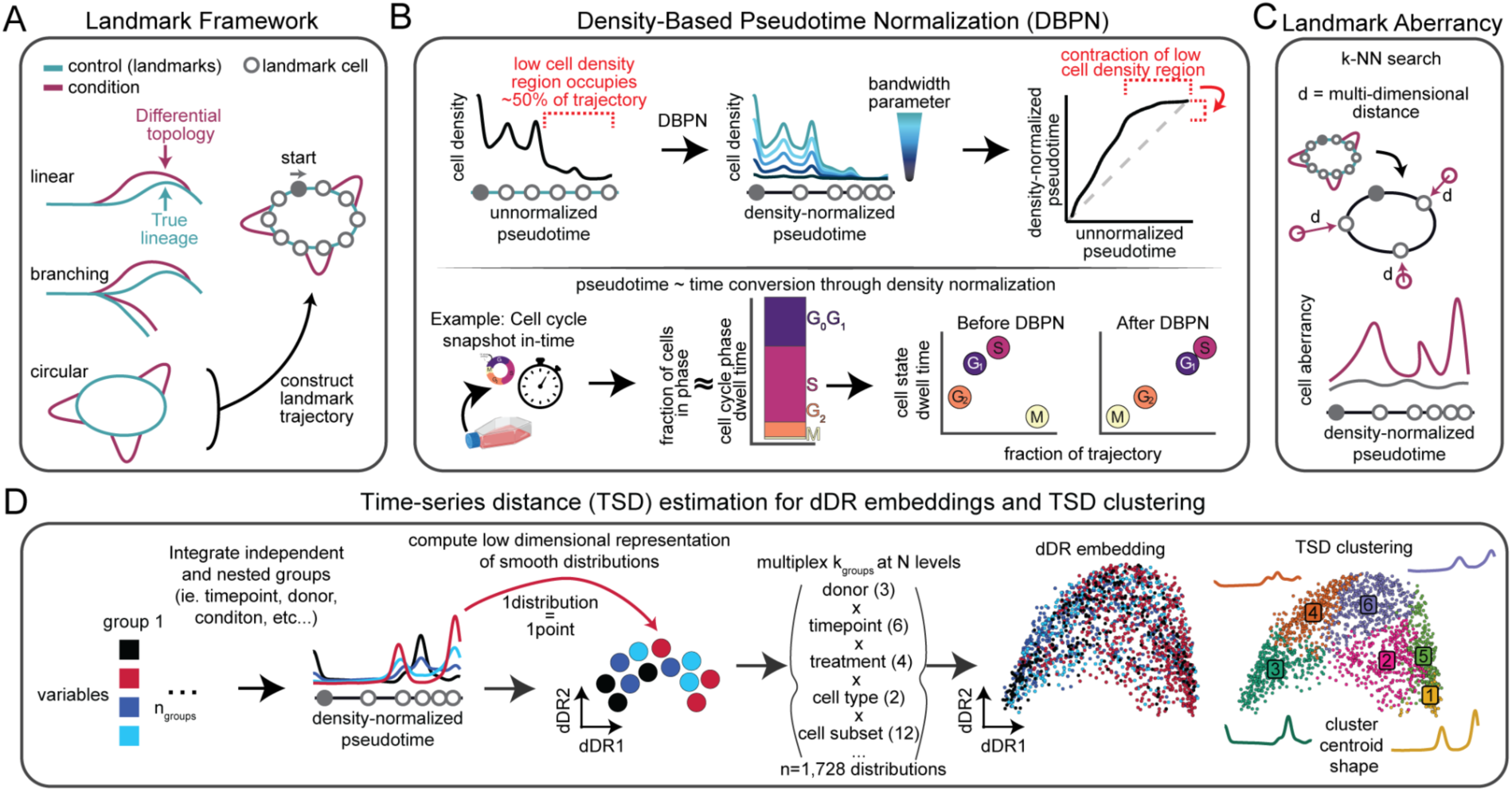
Hafez: a cell cycle landmark trajectory inference and time-series analysis framework generalizable to other systems. **(A)** Schematic of traditional single-cell lineage topologies and an example of a landmark-based trajectory inference strategy for cyclical topologies using the Hafez framework. **(B)** Cell density-based pseudotime normalization (DBPN) that expands and contracts pseudotime estimates (left). Pseudotime-time conversion after DBPN matches trajectory fraction with dwell time of cell states in a dynamic system (right). **(C)** Schematic of example landmark aberrancy analysis in Hafez using Euclidean distance of nearest neighbor landmark cells. **(D)** Schematic of Hafez’s time-series distance (TSD) estimation across different groups of interest for visualization using dDR^Hafez^ (dynamic time warping Dimensionality Reduction) embeddings and TSD clustering^Hafez^. Supplemental Figure 1 includes additional examples.

**Supplementary Figure 2:**
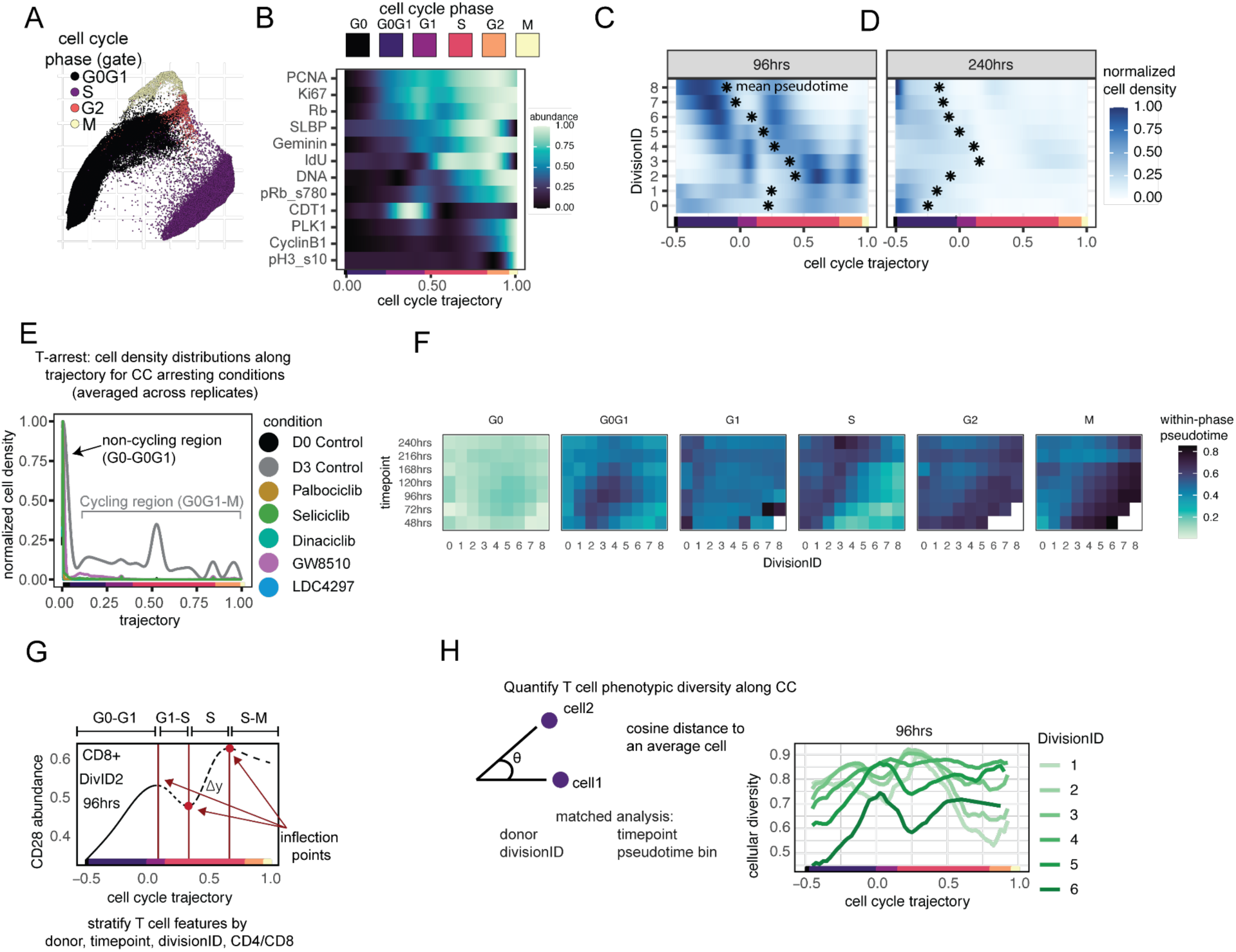
Cell cycle over time and division in primary human T cells. **(A)** T cell CC embedding across all donors colored with manually gated CC phases. **(B)** Molecular abundance of CC molecules along CC trajectory. **(C)** CC pseudotime across divisions at 96hrs and **(D)** 240 hrs, with points indicating mean pseudotime **(E)** Average cell density distributions along a cell cycle trajectory for each treatment group (D0 Control, n=1. D3 Control, n =3. Palbociclib, n=3. Seliciclib, n=3. Dinaciclib, n=3. GW8510, n=1. LDC4297, n=3). **(F)** Averaged CC cell density across all donors over time and division. **(F)** Average CC pseudotime across time and division normalized between 0 and 1 for each trajectory-annotated CC phase to highlight phases with potential time- or division-dependent changes in cell distributions. **(G)** Depiction of CD28 molecular abundance along a cell cycle trajectory with example inflection points. **(H)** Cosine similarity over divisions at 96hrs.

**Supplementary Figure 3:**
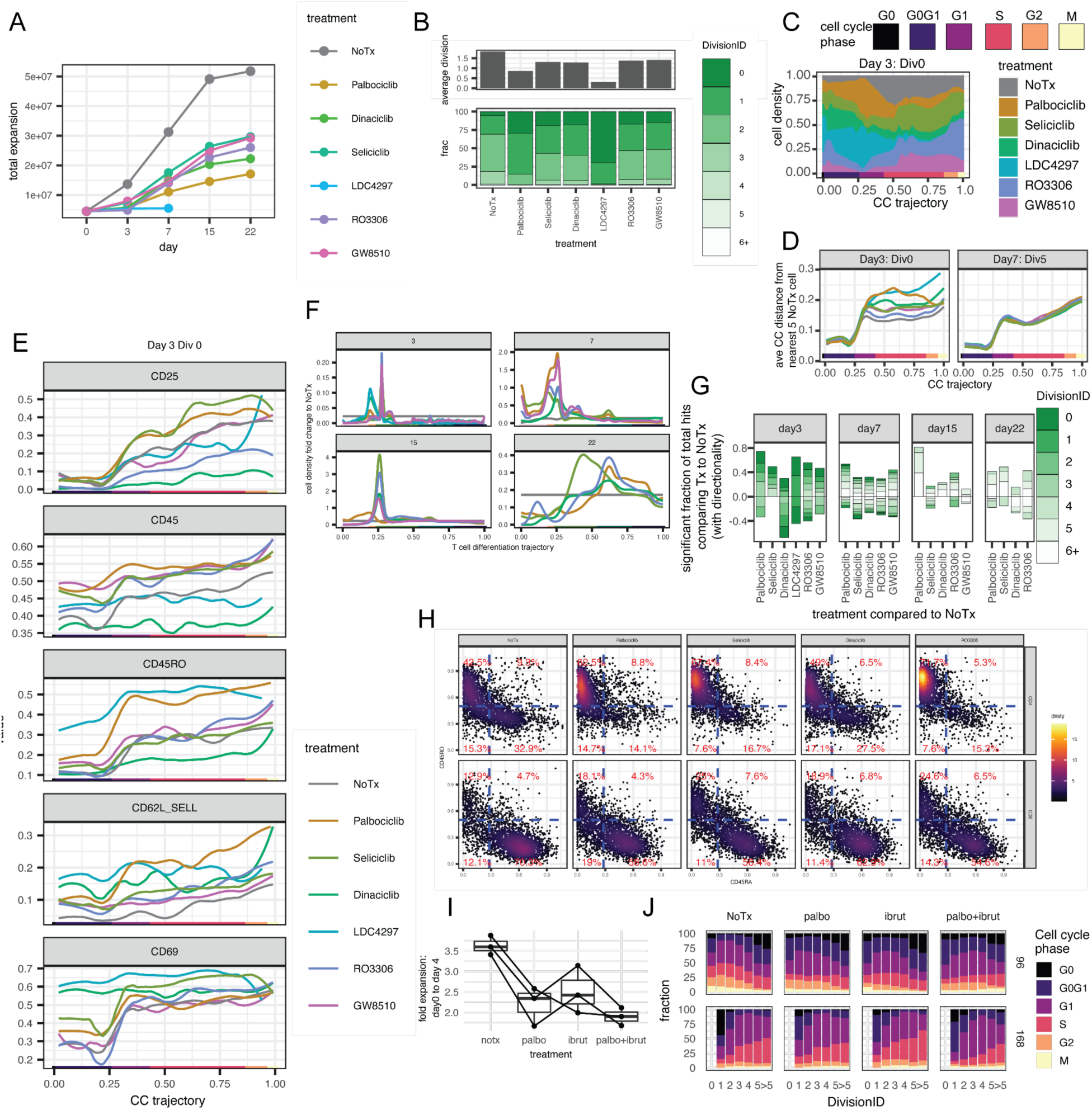
CDK inhibition and ITK blockade. **(A)** Viable cell expansion counts for each treatment over time. All drugs slowed reduced expansion over time. Cells treated with LDC4297 (inhibiting CDK7) did not survive after Day 7. **(B)** Average division based on CFSE annotations and the fraction of cells in each DIvision ID for day 3 post-stimulation. All drugs observed fewer divisions with treatment. **(C)** Normalized T cell density of all conditions along the CC trajectory constructed on untreated T cells. We observe more CC skewing towards G0, G0G1, and G1 states in undivided cells on day 3 compared to non-treatment. **(D)** NN aberrancy score computed using the Hafez framework. Drug-induced, early transient CC aberrancies in undivided cells on day 3 of drug washout resolved in highly divided cells by day 7. **(E)** Protein abundance of example markers for day 3, undivided cells. **(F)** Fold change in cell density for each treatment compared to Control within each timepoint. **(G)** Significant fractions summarized by Division IDs. **(H)** CD45RO vs CD45RA biaxial plot of day 22 cells with >5 divisions from each treatment split by CD4 and CD8 cells. **(I)** Fold expansion from day 0 to day 4 for each treatment group. **(J)** Trajectory-based CC annotations for each treatment over Division at 96hrs and 168hrs.

**Supplementary Figure 4:**
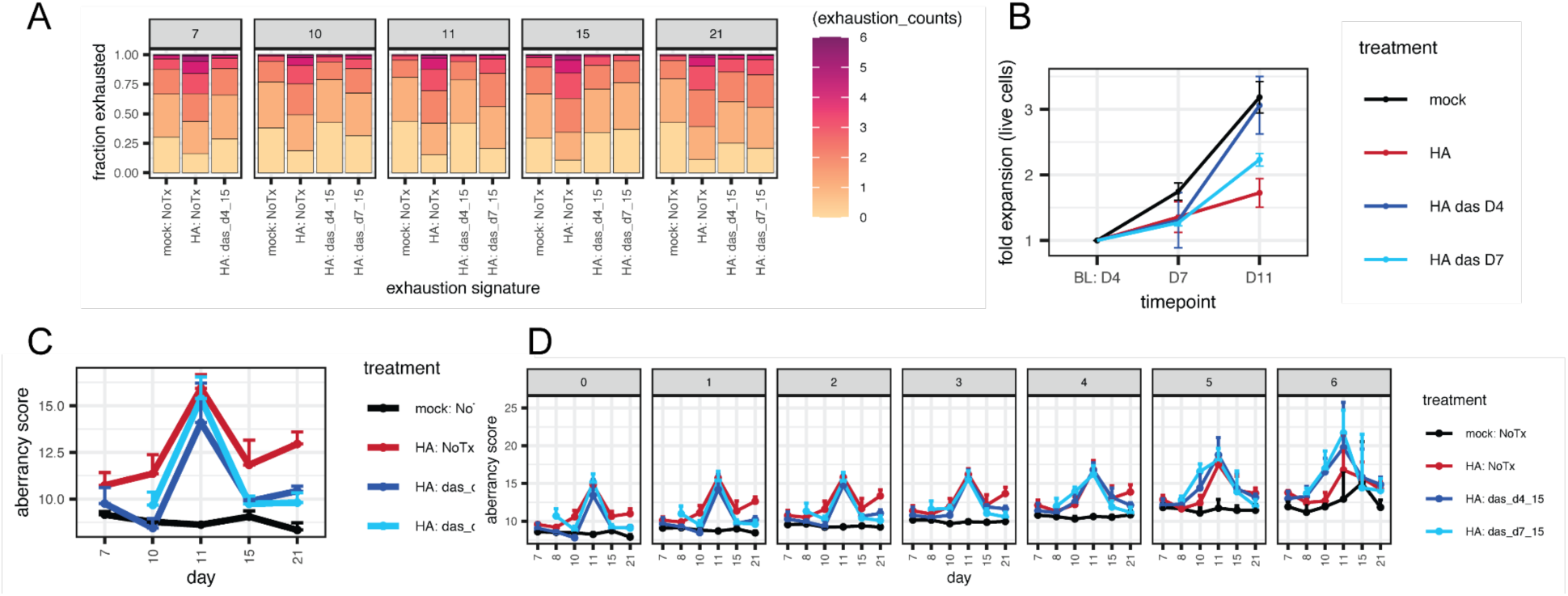
CC aberrancy analysis in Mock and HA CAR T cells treated with receptor signaling present or absent with dasatinib treatment. **(A)** Fraction of cells with exhaustion signatures. **(B)** CC aberrancy over time. **(C)** CC aberrancy over stratified by exhaustion score. **(D)** Differential features on each day for HA CAR T cell groups without and with dasatinib treatment compared to mock matched by donor and exhaustion signature.

**Supplementary Figure 5:**
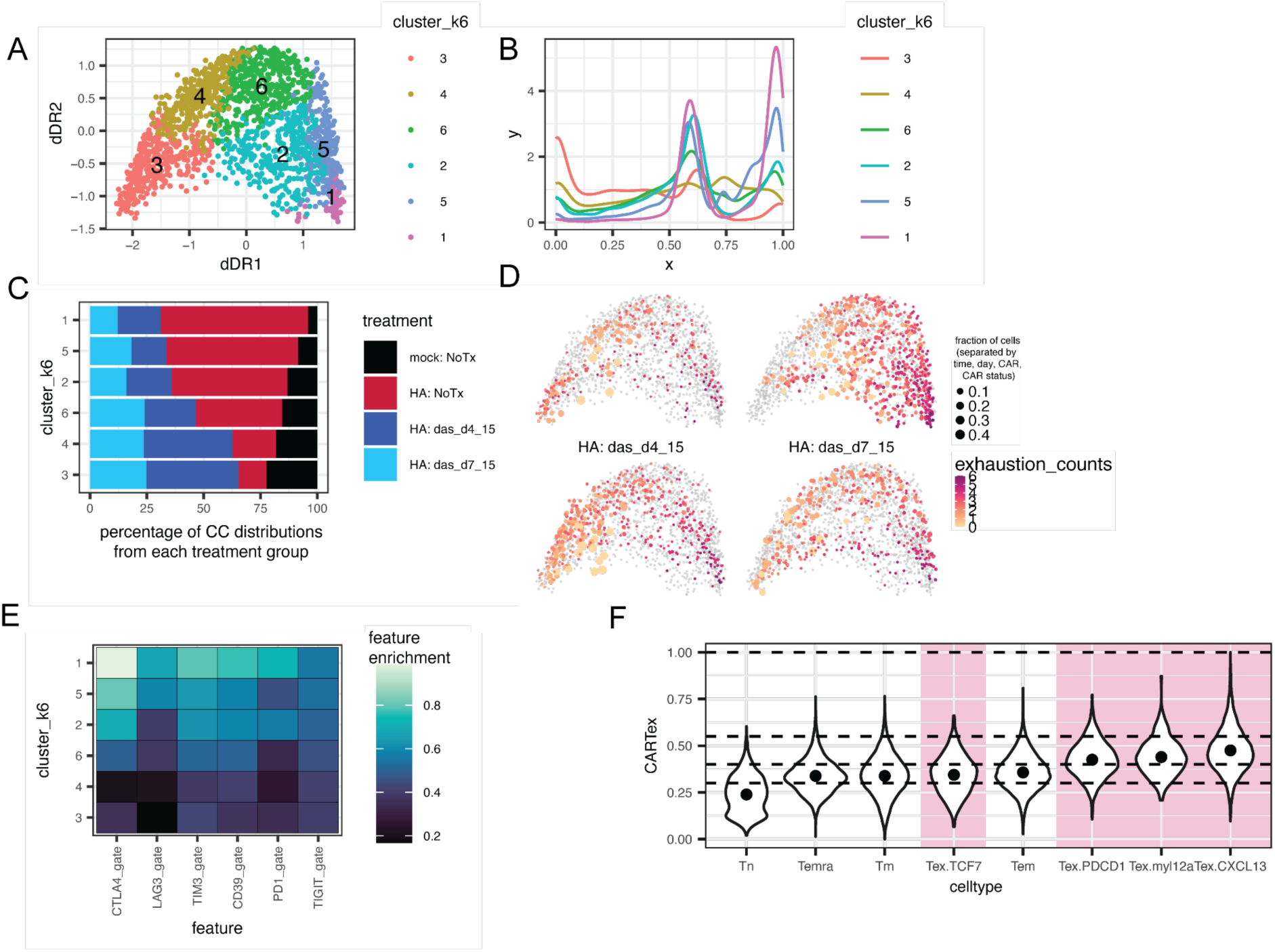
CC aberrancy in tonic signaling models of exhaustion and exhaustion scores in pan-cancer CD8 TILs. **(A)** *TSD* clusters of CC cell density patterns. **(B)** *TSD* clusters patterns. **(C)** Abundance of treatment groups in each *TSD* clusters (fractions). **(D)** dDR^Hafez^ embedding, with point size corresponding to the fraction of cells with each exhaustion signature. **(E)** Enrichment scores for each exhaustion marker in each cluster. **(F)** CARTex signature and CARTex bins for each celltype in the pan-cancer CD8 TILs dataset.

## Materials and methods

### Data availability

All data (fcs files) generated in this study are ready for public release upon publication. Any additional information desired in this article is available from the lead contact upon request.

### Code availability

Hafez software can be accessed on https://github.com/mamouzgar/hafez.

### Cell line culture

Five cell lines were used in this study: NALM6, U937, HEL, 293T (ATCC CRL-3216), JURKAT (ATCC CRL-2899). NALM6 is a lymphocyte-like cell line derived from human B cell precursor leukemia, U937 is a pro-monocytic cell line originating from human myeloid leukemia, HEL is an erythroblast cell line derived from human erythroleukemia, 293T is an adherent epithelial-like cell line originating from human embryonic kidney cells, and JURKAT is a CD4+ lymphocyte-like cell line derived from human T cell acute leukemia. The NALM6, JURKAT, U937, and HEL cells were maintained in RPMI 1640 (Gibco, 11-879-020), 10% FBS (Sigma-Aldrich, F4135), 1% p/s (Thermo Fisher Scientific, 15140122), and Glutamax (Thermo Fisher Scientific, 35-050-061). Cells were maintained at 37 °C, 5% CO2. 293T cells were maintained in DMEM (Dulbecco’s Modified Eagle’s Medium)/Hams F-12 50/50 Mix (Corning, 10-090-CV), 10% FBS, and 1% p/s.

### IdU labeling

5-Iodo-2-deoxyuridine (Sigma I7125) was re-suspended in DMSO (Sigma D2650) at 500 mM. IdU labeling was performed at a final concentration 100 μM and returned to the incubator for 15-30 minutes. After labeling, cells were washed with PBS and continued with the fixation protocol for CyTOF.

### Live/dead labeling, cell fixation, and permeabilization for CyTOF staining

CyTOF staining was performed as previously described, generally following this protocol.io publication (dx.doi.org/10.17504/protocols.io.14egn69nml5d/v2). Briefly, live-dead labeling was performed using cisplatin in low-barium PBS, cells were washed with CSM, fixed with PFA for 10 minutes at room temperature, and stored in -80C until staining. When ready, fixed samples were thawed at room temperature and barcoded using fixed-cell palladium barcoding, pooled into a single tube. Surface staining was performed with metal-conjugated antibodies in CSM for 30 minutes at room temperature, washed, permeabilized with 100% 4c methanol and incubated on ice for 10 minutes, washed 3x times with CSM, and proceeded with intracellular staining. Finally, cells were washed with CSM, and resuspended in an intercalator solution until CyTOF analysis.

### Palladium barcoding and staining with metal-conjugated antibodies

Individual samples within one experiment were palladium barcoded as described previously (https://doi.org/10.1038/nprot.2015.020) and combined into a single sample before further processing and staining. Experiments with multiple barcode plates had an anchor sample to normalize any technical effects.

### Antibody and molecular targets

Supplementary Table S1 contains a detailed list of all molecular targets used or measured, including metal-conjugated cell cycle and T cell targets, and biotinylated antibody target clones along with their vendors, concentrations, and metal channels when applicable.

### Ex vivo Primary human T cell isolation, culture, and TCR stimulation

Deidentified human blood was obtained from healthy adult donors below 40 years old under informed consent. Bulk T cells were negatively isolated from whole blood using RosetteSep Human T Cell Enrichment Cocktail (StemCell Technologies). When applicable, untouched Naive T cells were negatively isolated using using Streptavidin Particles Plus (BD Biosciences #557812) and the BD IMag Cell Separation Magnet (BD Biosciences) with biotinylated anti-CD45RO, anti-CD57, and anti-CD95 (**Supplementary Table S1**). T cells were cultured in ImmunoCult-XF T Cell Expansion Medium (10981, Stemcell Technologies) and supplemented with 10 ng ml−1 of interleukin-2 (Miltenyi Biotec). Briefly, cells were incubated for 15 minutes on ice using Fc block, the antibody cocktail was added (1 ug/5million cells) and cells were incubated for 15 minutes on ice, washed with PBS+0.5% BSA and EDTA, streptavidin particles were added at 50 uL/10e7 cells and incubated for 30 minutes at room temperature before transfer to the BD Imag separation magnet for 8 minutes to collect untouched naive T cells with 3 rounds of separation to increase purity. Bulk or native T cells were activated and expanded using Human T-Expander CD3/CD28 (Dynabeads, Thermo Fisher) added in a 1:1 cell-to-bead ratio and cells were incubated at 37 °C in 5% CO2. Where applicable, naive T cells were negatively isolated using biotinylated CD45RO, CD57, and CD95 to obtain untouched Tn cells. CFSE incorporation was performed by incubating cells with 80 μm of CFSE (Thermo Fisher) in CCM for 5 min at room temperature (RT) as described previously (https://doi.org/10.1038/s41587-019-0033-2). Labeled cells were quenched with warm CCM and washed three times by centrifuging for 5 min at 200g. T cells were cultured in ImmunoCult-XF T Cell Expansion Medium (10981, Stemcell Technologies) and supplemented with 10 ng ml−1 of interleukin-2 (Miltenyi Biotec). T cells were activated and expanded using Human T-Expander CD3/CD28 (Dynabeads, Thermo Fisher) added in a 1:1 cell-to-bead ratio and cells were incubated at 37 °C in 5% CO2.

### Drug treatments

Drug treatments were performed using ibrutinib (PCI-32765; Cellagen Technology #C7327), palbociclib (Chemscene, CS-3110), roscovitine (Seliciclib, CYC202, Selleck Chemicals #S1153), dinaciclib (SCH727965, Selleck Chemicals, #S2768), LDC4297 (LDC044297, Selleck Chemicals, #S7992), ro3306 (Selleck Chemicals,#7747), GW6510 (sc-biotech #sc-215122), hydroxyurea (Sigma-Aldrich #H8627-1G), nocodazole (Sigma-Aldrich. #SML1665-1ML), SU9516 (EMD Millipore, Calbiochem, #572650), and DMSO (Sigma-Aldrich D2650) for untreated/vehicle. Concentrations are provided in the Supplementary Tables. Freshly thawed dasatinib (1um) was mixed with T cell culture medium and supplemented every 2 to 3 days.

### CyTOF processing

Raw mass cytometry data were bead normalized to remove acquisition-related influences on marker expression using the premessa R package. Sample barcoding was done using fixed cell palladium barcode combinations and debarcoded using premessa. Normalized data were uploaded to CellEngine for bead removal, singlet identification, removal of debris and non-biological events, live cell gating, and exported (https://www.cellengine.com/). Pre-apoptotic cells defined by cPARP positivity were also removed. Batch correction between multiple palladium barcodes in the same experiment was performed using an anchor sample with an adjustment factor for each channel (https://doi.org/10.3389/fimmu.2019.02367). Adjusted data was imported into CellEngine for manual cell cycle gating using CyclinB1, IdU labeling, and phospho-HistoneH3 (s10). Gated files were subsequently imported into the R environment, asinh transformed (cofactor 5), and normalized to the 99.9th percentile of each respective channel before downstream analysis.

### Application of Density-based Pseudotime Normalization (DBPN)

For method details, please see the method section subtitle for Hafez Framework. DBPN for analysis applications was used on control cells unless otherwise noted (untreated, healthy donor). In multi-donor and multi-timepoint analysis experiments, we equally sampled cells from each donor and a range of timepoints to capture the breadth of cell cycle states (i.e, Day 0 unactivated and a series of later timepoints). For normalizing trajectories with respect to drug treatments, DBPN was validated experimentally using cell cycle inhibitors that arrest the cells, increasing their dwell-time in specific cell cycle states. In this setting, trajectories are computed on control cells and all other cells are projected. Instead of using untreated cells as the input into DBPN, we input the drug-treated cells and recompute the pseudotime normalization for each drug separately using the same original cell ranking to remap the pseudotime distribution according to the inhibitor action.

### Dimensionality reduction

Reduced embeddings were constructed using either PHATE (R/Python) with knn_dist=cosine, mds_dist=euclidean, and knn=15 or uwot::UMAP (R) using default parameters. We used RANN::nn2 in R or sklearn’s KNeighborsClassifier in Python to find the nearest WT or untreated cell neighbors by euclidean or cosine distance. Analysis and plotting was done in R v4.2.

### Interaction analysis

Interaction analysis of single order, two-order, three-order, and four-order combinations of different T cell subsets was performed by calculating the enrichment score for different combinations using meld with 16 T cell targets (2,516 total combinations). The top 30 strongest combinations were extracted and the sum of the scores for each interacting pair was calculated and plotted using ggraph and tidygraph.

### Single-cell RNAseq data curation and analysis

Single-cell RNAseq data was obtained from published GSE repositories. The const and snp CAR T cells were downloaded from GSE192998, and the pan-cancer TILs were downloaded from GSE156728. Pre-processed data was analyzed using Seurat. Const/snp CAR T cells were annotated with cell cycle phases using Seurat’s CellCycleScoring function, and RNA velocity analysis was performed using scVelo. Pan-cancer TILs were cell cycle scored and a cell cycle ordering computed using methods in Amouzgar et al, 2022.

### Statistical analysis

Generalized linear model (glm) and generalized linear mixed model (glmm) as used in the diffcyt framework was used for statistical analysis of CyTOF data. All statistics were performed using matched analysis between donors, and additional covariates were included when relevant (timepoint, CAR type, exhaustion score, DivisionID, etc). When multiple comparison groups were compared to each other (eg, multiple treatment groups), we used estimated marginal means procedure to identify statistically different associations between groups of interest.

Multiple hypothesis correction was performed across all comparisons and features of interest in an experiment using the bonferroni correction and significance was determined by an adjusted p-value <= 0.10, unless otherwise noted. When noted, a fold-change/coefficient estimate threshold was also included to determine different features of interest between comparison groups. To avoid inflated p-values from single-cell observations, pseudobulk estimates were computed (eg, mean expression in each donor, timepoint, DivisionID). For statistically comparing CC distributions to exhaustion counts, we quantified the spearman correlation between exhaustion counts and the weighted mean value of the CC distribution. When relevant, CC distributions were statistically compared (eg CD19 vs HA) by computing the absolute difference (observed test statistic) between the groups of interest, randomly permuting the group labels 1000 times, and then determining whether the observed mean is greater than the permuted test statistic’s distribution. For Supplementary Figure 5, lasso regression was used to predict CC pseudotime from interaction terms between time and division within each CC phase as a means to identify whether there were intra-phase changes in cell density enrichment as a consequence of time or division. Generalized additive models (GAM) were computed using penalized regression splines in the mgcv::gam package in R to compute smoothed lines. Statistical analysis is provided as supplementary tables.

### Hafez framework

The name, Hafez, is inspired by the 14th century Persian poet whose collected works are regarded as some of the greatest achievements in Iranian literature. It is tradition to delve into the mysteries of fate and destiny through “faal-e-Hafez’’ (divination) when faced with challenges, choice, questions, or for entertainment. Hafez’s poetry intends to reveal our fate or destiny during transitions in life, drawing parallels to the field of single-cell trajectory inference that maps cell fate and reveals cell state transitions. The word Hafez literally translates to “one who remembers” or “keeps in memory” - which is fitting considering cell fate mechanisms are often encoded earlier in differentiation and remembered as cells meet their destiny. Hafez often ponders the interconnectedness of human actions (experimental) and cosmic forces (computational) to interpret the world, so we pay homage to his legacy by naming our integrated experimental and computational method for interpretable landmark trajectory inference and systems modeling, Hafez.

Standard TI strategies discover lineages on single cells from all conditions. With single-cell technologies enabling greater multiplexing of disease or perturbation conditions with diverse differential topologies, current TI strategies may encounter difficulties capturing human-interpretable lineages if the scale of aberrant cell states from multiple disease or perturbation conditions overwhelms the presence of healthy, normal cell states^12^. Solutions to this include the use of dynamic programming strategies such as dynamic time warping to optimally map timepoints along trajectories (CellAlign)^12,62,63^, reference and query mapping at single-feature resolutions to realign trajectories (Genes2Genes)^63^, and mapping procedures that embed new cells onto reference embeddings^17^. However, these algorithms are not benchmarked for other single-cell technologies such as CyTOF, which is uniquely suitable for deep CC analysis because of its generalizability to suspension, adherent, and dissociated cell cultures, sample multiplexing, and parallel measurement of protein, phospho-protein, and DNA labeling strategies^26^.

The Hafez framework integrates multiple analysis strategies for trajectory inference and post-inference pseudotime analysis, and is designed using cell cycle biology as a basis - though we demonstrate the utility of Hafez in analyzing other biological systems as well. This framework primarily consists of three features: (1) landmark-based trajectory inference to purposefully capture an interpretable trajectory grounded in known biology with a normalization and projection approach to embed out-of-sample cells onto the landmark trajectory; (2) density-based pseudotime normalization (DBPN) to remap the pseudotime estimates of cell states to the dwell-time of cell states; (3) Post-trajectory analysis architecture that can leverage the landmark trajectory framework along a shared trajectory. *Hafez is considered an additional toolkit for trajectory inference. It is not designed to replace existing trajectory inference strategies and its performance as a trajectory inference algorithm is not meant to be better or worse than other strategies, but situational. Thus, users can use Hafez analysis strategies with other trajectory inference algorithms.* That said, we provide substantial benchmarking across supplemental figures 2-4 to demonstrate the utility of a landmark framework and its downstream functions across different datasets. We quantify and benchmark this strategy compared to the consequences of different sampling strategies for TI using 5 different datasets: CC inhibitors in (i) primary human T cells (main text), (ii) multiple cell lines (Figure S3B-D), (iii) jurkat cells with drug treatments (Figure S3F, S4A-D), (iv) *ex vivo* naive CD8^+^ T cell stimulation measuring the metabolic regulome across multiple timepoints (Figure S3G-I), and (v) B cell development in healthy and BCP-ALL (Figure S4E-M).

1. *Landmark trajectory inference:* Elastic principal graphs are used as the foundation for the landmark-based trajectory inference strategy^64,65^. After constructing the graph structure using the landmark cells, out-of-sample test data can be projected onto the graph architecture using our customized projection approach where landmark cells can be optionally defined for preprocessing and extension to the test data.
2. DBPN: DBPN for trajectory remapping to an unperturbed or other landmark cell system’s distribution can improve trajectory interpretability by mapping CC pseudotime to known CC phase lengths, metabolic state, or other cell state durations. In any TI setting, it may be valuable to perform DBPN on healthy or untreated samples to assist in trajectory interpretation through the lens of normal biological progression or using any other set of landmark cells most sensible for downstream analysis. Data normalization and processing strategies like PCA transforms the data, and the selection of reduced dimensions can influence the trajectory inference output. A consequence of different data processing strategies is that the cell density along the pseudotime of a trajectory inference analysis may also change even if the general rank-order of the cells along a trajectory might not differ and the transition states are accurately captured. DBPN uses a kernel density estimation approach to smoothly expand and contract the pseudotime based on cell abundance at a pseudotime-point in the trajectory, effectively mapping the pseudotime range to the dwell-time of cell states for either all cells or a desired set of cells (eg, landmark cells), grounding the cell states along the pseudotime to the dwell-time biology of the system (see main figures and supplement for experimental validation). DBPN improves pseudotime interpretability by shrinking the pseudotime occupancy of rare cell states that might occupy a large fraction of a calculated pseudotime (eg, mitotic cells in a cell cycle trajectory). This is particularly useful when perturbation or aberrant cell systems are studied as described in the figures because the pseudotime can be normalized to healthy or untreated landmark cells, and other conditions can be visualized in reference to the landmark trajectory. Cell density is calculated using a univariate gaussian kernel density estimation given as positive function p*K*(y; bw) which is controlled by the bandwidth (bw) parameter. The density estimate at point *y* within a group of pseudotime bins *xi*; *i*=1 … *N*:

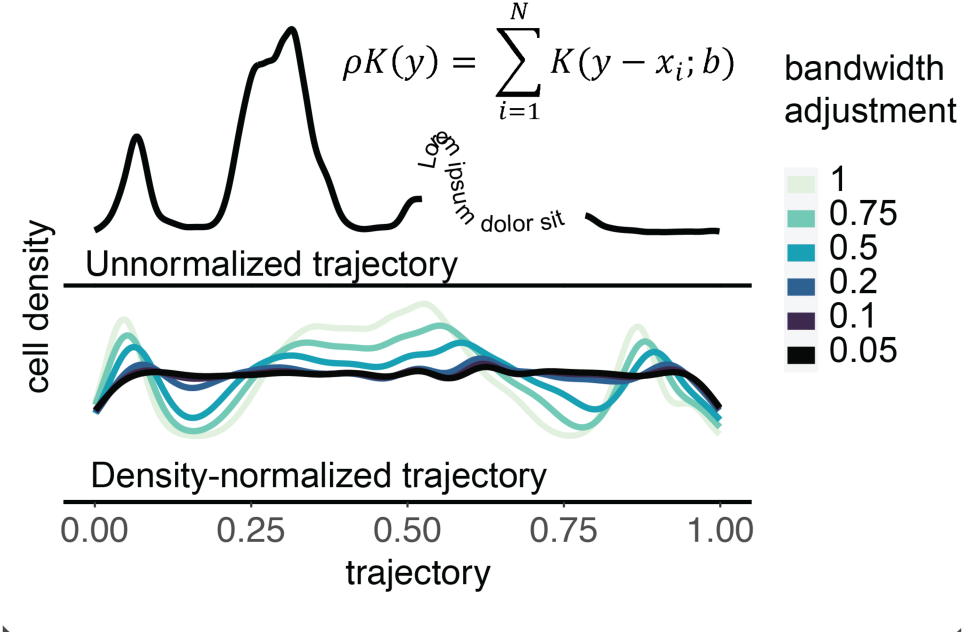 Large bandwidth values lead to very smooth density distributions, capturing fewer nuances in the dwell-time of cell states while small bandwidth values capture more granular nuances in dwell-time cell states. The density estimation for each pseudotime bin is divided by the sum of all density points, and the new start point for each pseudotime bin is calculated using the cumulative sum. A lag operator is used to add the prior bin’s cumulative sum to the minimum bin start point before rescaling the pseudotime values within the bin. This effectively expands and contracts the pseudotime, normalizing the pseudotime trajectory to match the dwell-time of cell states. In the case where landmark cells are used as the reference set to perform the normalization, the remaining test cells are mapped onto the newly-normalized pseudotime using a simple SQL join query before rescaling the pseudotime according to the landmark-computed density estimation. To equalize for different cell estimates in multi-sample experiments, DBPN can optionally calculate the density estimation for each sample separately and then average the density distributions before executing the normalization procedure.
3. Post-trajectory, time-series analysis architecture: Hafez includes multiple methods for post-trajectory analysis to analyze data that can leverage the landmark cell framework. However, these are just examples and we invite others to use the landmark cell framework for other *a*nalysis strategies. This includes landmark-based cell density estimation, multivariate comparison of conditions using pseudotime-mapped mahalanobis distance to landmark cells, dimensionality reduction of time-series distances (TSD), and TSD clustering of trajectory patterns.

a. Landmark-based cell density estimation: Cell density estimation is calculated using the same univariate kernel density estimation approach as used in DBPN, excluding the pseudotime normalization procedure. Density estimation is separately computed for each group of interest, *f^g^*, using the same user-specified bandwidth value.
b. Pseudotime-mapped scores: Pseudotime discretized and different metrics can be used as desired, including mahalanobis or euclidean metrics as aberrancy scores, or cosine distance. Mean and covariance matrices are calculated for each pseudotime bin using ether (a) all cells in a pseudotime bin and/or (b) user-specified landmark cells.
c. Time-series distance (TSD) visualization and clustering for pattern identification using dynamic time warping. Deeper analysis of pseudotime patterns independent of magnitude can be performed by cell density estimations or smoothed molecular programs (as in this paper) by different groups. TSD is computed using dynamic time warping to compute distances followed by time series clustering using dtwclust^66^.

## Acknowledgements

We thank Angie Spence and Mako Goldston for their help with antibody conjugations. Figures were plotted in R, cartoons generated using BioRender (http://www.biorender.com) Adobe Illustrator.

## Contributions

MA and SCB conceived and conceptualized the study. MA conceived, conceptualized, and wrote all analysis methods, software, and code. MA performed cell line, T cell culture, perturbations, mass cytometry experiments, validated reagents, analyzed all the data, and wrote the manuscript. PF and AJL helped design and perform cell culture experiments, and edited the manuscript. TM helped with CAR T cell transductions and cultures. TM, ES, and CM helped with conceptualization of CAR T cell experiments. TB and DH conjugated metal-tagged antibodies, helped with both antibody validation and mass cytometry experiments, and negatively isolated bulk T cells. S.C.B. supervised the study, provided resources, acquired funding, wrote and edited the manuscript.

